# Single-cell Individual Complete mtDNA Sequencing Uncovers Hidden Mitochondrial Heterogeneity in Human and Mouse Oocytes

**DOI:** 10.1101/2020.12.28.424537

**Authors:** Chongwei Bi, Lin Wang, Yong Fan, Gerardo Ramos-Mandujano, Baolei Yuan, Xuan Zhou, Jincheng Wang, Yanjiao Shao, Pu-Yao Zhang, Yanyi Huang, Yang Yu, Juan Carlos Izpisua Belmonte, Mo Li

## Abstract

The ontogeny and dynamics of mtDNA heteroplasmy remain unclear due to limitations of current mtDNA sequencing methods. We developed individual <u>Mi</u>tochondrial <u>G</u>enome <u>seq</u>uencing (iMiGseq) of full-length mtDNA for ultra-sensitive variant detection, complete haplotyping, and unbiased evaluation of heteroplasmy levels, all at the individual mtDNA molecule level. iMiGseq uncovers unappreciated levels of heteroplasmic variants in single healthy human oocytes well below the current 1% detection limit, of which numerous variants are detrimental and could contribute to late-onset mitochondrial disease and cancer. Extreme mtDNA heterogeneity among oocytes of the same mouse female, and a strong selection against deleterious mutations in human oocytes are observed. iMiGseq could comprehensively characterize and haplotype single-nucleotide and structural variants of mtDNA and their genetic linkage in NARP/Leigh syndrome patient-derived cells. Therefore, iMiGseq could not only elucidate the mitochondrial etiology of diseases, but also help diagnose and prevent mitochondrial diseases with unprecedented precision.

## Introduction

Mitochondria play vital roles in cellular metabolic and signaling processes. Each human mitochondrion contains on average 1.4 copies of a 16.5 kb circular genome^1^–mtDNA– that is densely packed with 13 genes encoding core subunits of the oxidative phosphorylation complexes, 24 RNA genes, and a non-coding control region (D-loop region). Mitochondria and hence mtDNA undergo constant turnover even in non-dividing cells such as arrested primary oocytes^2, 3^. The demand for frequent DNA replication and the lack of histonized chromatin in mtDNA contribute to a mutation rate that is at least one order of magnitude higher than that of the human nuclear genome^3–5^. Because mammalian cells typically contain 1,000-10,000 copies of mtDNA, mutations arisen in individual mtDNA produce an admixture of mutant and wild-type mtDNA (heteroplasmy) within a cell^6^. Heteroplasmic mutations, inherited exclusively from the oocyte, can cause inborn disorders and are associated with late-onset complex diseases^4^.

Our understanding of heteroplasmy has been dramatically shaped by available technologies, from being perceived as a rare phenomenon in early studies to one that affects nearly 1 in 2 individuals^7, 8^. In recent years, deep short-read next-generation sequencing (NGS) based methods have been instrumental in revealing the rich diversity of mtDNA. They were used to show that heteroplasmic mutations are present in the general population and are among the most common causes of inherited metabolic diseases when present above a heteroplasmy threshold^9, 10^. However, previous findings are susceptible to the limitations and biases of existing sequencing strategies. The reported heteroplasmic levels are inferred from sequencing of PCR products amplified from a heterogeneous population of mtDNA, and thus can be inaccurate due to PCR bias^11^. Most importantly, current NGS-based studies of mtDNA are blind to rare heteroplasmic variants that exist below 1% allele frequency due to the background error rate (0.1∼1%) of NGS and contamination of nuclear mitochondrial DNA-like sequences (NUMTs) in short amplicon PCR^3, 12, 13^. As such, the prevalence, nature, and significance of these rare heteroplasmic variants remain unknown.

Another challenge of the field is the lack of quantitative analysis of the multitude of mitochondrial genomes at the individual-molecule level in single cells, which would be necessary to study the ontogeny, heterogeneity, dynamics, and genotype-phenotype relationship of mtDNA mutations. Current methods of mtDNA sequencing suffer from two seemingly paradoxical issues–amalgamation and fragmentation. Most studies of mtDNA genetics are based on short-read shotgun or amplicon next-generation sequencing (NGS)^2, 12–16^. These techniques average out the heterogeneity of mtDNA in two ways. Firstly, mtDNA genotypes of thousands of cells are averaged in bulk sequencing, thus masking variants in rare cells and cell-to-cell heterogeneity. Secondly, even in single-cell analysis^17, 18^, it was a composite genotype of all mtDNA rather than the true genotypes of individual mtDNA that was obtained. Additionally, the phenotypic significance of an mtDNA variant is strongly modified by other co-inherited variants^4^. However, conventional methods cannot provide full haplotypes due to fragmentation of mtDNA molecules, which largely prevents the study of linkage between heteroplasmy variants.

For the afore-mentioned reasons, high-throughput analysis of mutations in a single mtDNA molecule in single cells is still beyond the reach of current methodologies. To overcome these hurdles, we applied a long-read individual molecule sequencing strategy (IDMseq^19^), which is several orders of magnitude more sensitive (capable of detecting allele frequency as low as 0.004%) than conventional NGS and provides haplotype-resolved quantitative analysis of variants, to sequence individual complete mtDNA in single oocytes. This new technology– individual <u>Mi</u>tochondrial <u>G</u>enome <u>seq</u>uencing (iMiGseq)–labels each mtDNA in single cells with a unique molecular identifier (UMI). UMI-labeled mtDNA are further amplified by high-fidelity long-range PCR and sequenced on long-read sequencing platforms to obtain full-length mtDNA, generate variant details, and assemble the genome for individual mtDNA (Fig. 1a).

**Fig. 1.**
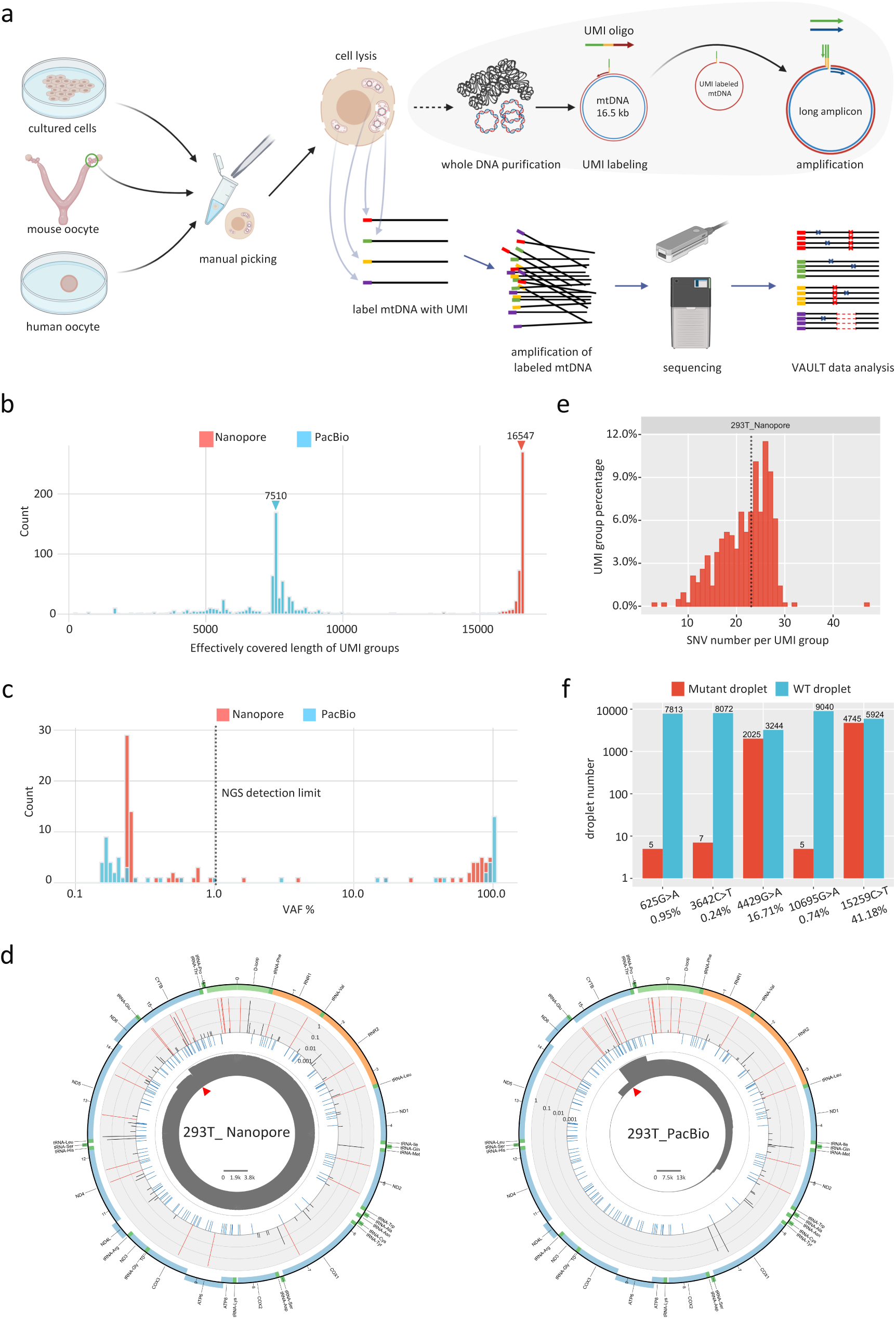
Validation of iMiGseq using 293T cells. **a**, Schematic representation of iMiGseq. Single cells (5 cells for 293T) were manually picked and lysed in RIPA buffer. iMiGseq applies the IDMseq strategy to specifically label individual mtDNA with UMIs by single-oligo priming. The UMI oligo contains a 3’ gene-specific sequence (red), a UMI sequence (yellow), and a 5’ universal primer sequence (green). The UMI-labeled mtDNA is further amplified by long-range PCR using the universal primer (green) and gene-specific reverse primer (purple) as a single amplicon. The sequencing of long amplicons is performed on long-read Nanopore and PacBio sequencing platforms. The sequencing data were analyzed by a bioinformatic toolkit–VAULT–to identify UMI sequences, bin reads based on UMI and call variants. **b**, Distribution of effectively covered length of UMI groups detected by Nanopore and PacBio sequencing. Color-coded triangles indicate N50 values. **c**, VAF distribution of SNVs detected in iMiGseq of 293T cells. The majority of unique SNVs are below the current 1% detection limit (the vertical dotted line). **d**, Circular plots showing the distribution of SNVs in the mitochondrial genome of 293T cells determined by Nanopore and PacBio sequencing. The innermost circle (grey) shows the depth of reads of all detected UMI groups in linear scale as indicated by the scale bar in the center. The red triangle indicates the position of the primers. The middle circle (light blue) represents common SNPs from the human dbSNP-151 database. Individual SNVs are plotted in the outer barplot circle, in which the height of bar represents the VAF. Red color indicates VAF > 0.6. The outermost circle is a color-coded diagram of the human mtDNA. Blue: protein-coding genes. Yellow: rRNA genes. Dark green: tRNA genes. Light green: D-loop. **e**, Histogram of SNV load per mtDNA in 293T cells based on Nanopore iMiGseq. The median SNV number is indicated by the vertical dotted line. **f**, ddPCR results showing the detected events (droplets) for variants with different VAFs. The VAFs calculated by Nanopore iMiGseq are shown under the variants. Around 80 cells were used in ddPCR to ensure the generation of enough events for analysis, as compared to 5 cells in iMiGseq.

iMiGseq allowed us to address several key open questions in the field. It provided true heteroplasmy levels without the influence of PCR biases and NUMT contaminations. It uncovered pathogenic mtDNA mutations in human oocytes that lie below the current 1% detection limit. It showed the first haplotype-resolved mitochondrial genomes from single human oocytes. It revealed the linkage of rare heteroplasmic mutations and allowed the study of the linkage between mtDNA mutations. It also provided strong evidence of selection imposed on mtDNA variants during human germline transmission. Lastly, we showed that iMiGseq could characterized single-nucleotide variants (SNVs) and structural variants (SVs) of mtDNA in cells of a mitochondrial disease patient, thus facilitating diagnosis and prevention of mitochondrial diseases.

## Results

### Validating iMiGseq of full-length mtDNA

The ideal sequencing technology for iMiGseq should offer accurate ultra-long reads (∼16.5 kb). We have previously showed that the key components of iMiGseq, IDMseq and VAULT, have been validated to provide faithful quantitative characterization of various types of variants with frequencies as low as 0.004% through error correction by the molecular consensus strategy^19^. The design of iMiGseq further eliminated the influence of NUMT contaminations that perplexed the current NGS-based analysis of mtDNA variants (see methods).

We first tested two state-of-the-art long-read platforms–Oxford Nanopore MinION and PacBio Sequel. Five HEK 293T cells (∼1.5k copies of mtDNA/cell) were subjected to UMI labeling of full-length mtDNA (Fig. 1a, Supplementary Fig. 1a & b). Nanopore sequencing generated 81.8k reads mapped to the human reference mtDNA with an average alignment identity of 89.9% (Table 1, Supplementary Fig. 2a). Using the VAULT pipeline^19^, the reads were assigned to 542 UMI groups (a set of reads sharing the same UMI), of which 92.6% covered ≥ 95% of the full-length mtDNA (position with depth ≥ 3) and the N50 was 16,547 bp (Fig. 1b). After further filtering based on in-group read consistency, we obtained 426 high-confidence UMI groups, of which 390 covered ≥ 95% of the mtDNA, which represented 390 individual complete mitochondrial genomes. Variant analysis showed that 426 UMI groups contained a total of 8,986 high-confidence SNVs. No SVs were detected.

**Table 1.** Summary of individual sequencing runs.

| Sample | Mapped Read count | Reads with UMI | Detected mtDNA (with $\geq 5$ reads) | mtDNA with SNV | SNV count | Median covered region of UMI groups | high coverage mtDNA* | Average SNV load per mtDNA* | unique SNV number | somatic SNV (<0.01 VAF)* | somatic SNV load per mb* |
| --- | --- | --- | --- | --- | --- | --- | --- | --- | --- | --- | --- |
| 293T Nanopore | 81,798 | 21,635 | 426 | 426 | 8,986 | 16,546 | 390 | 21.72 | 88 | 64 | 9.9 |
| 293T PacBio | 166,194 | 51,096 | 637 | 637 | 9,368 | 7,510 | 0 | N/A | 57 | N/A | N/A |
| NSG mouse oocyte 1 | 1,086,723 | 245,363 | 4,078 | 175 | 182 | 10,494 | 1,322 | 0.05 | 85 | 47 | 2.2 |
| NSG mouse oocyte 2 | 959,528 | 193,793 | 1,165 | 131 | 134 | 15,704 | 596 | 0.13 | 44 | 36 | 3.7 |
| NSG mouse oocyte 3 | 1,992,516 | 612,227 | 19,697 | 225 | 237 | 5,562 | 4,384 | 0.03 | 98 | 117 | 1.7 |
| B6 mouse oocyte 1 | 988,865 | 200,174 | 2,493 | 1,173 | 2,145 | 7,782 | 436 | 2.44 | 44 | 14 | 2.0 |
| B6 mouse oocyte 2 | 549,736 | 116,430 | 1,849 | 945 | 1,818 | 8,573 | 363 | 2.57 | 40 | 13 | 2.2 |
| B6 mouse oocyte 3 | 325,899 | 63,039 | 615 | 329 | 557 | 9,926 | 129 | 1.97 | 14 | 2 | 1.0 |
| Human oocyte 1 | 763,582 | 95,387 | 2,308 | 2,301 | 75,483 | 16,531 | 2,072 | 34.80 | 141 | 128 | 3.7 |
| Human oocyte 2 | 397,426 | 64,646 | 1,461 | 1,457 | 32,276 | 16,413 | 1,009 | 27.30 | 81 | 59 | 3.6 |
| Human oocyte 3 | 506,082 | 99,404 | 1,535 | 1,514 | 22,522 | 15,568 | 712 | 23 | 69 | 20 | 1.7 |
| hEPs<sup>m.8993T>C</sup> | 72,496 | 34,959 | 536 | 536 | 12,173 | 16,573 | 515 | 23.19 | 117 | 72 | 8.5 |

\* data analyzed using extensive-coverage UMI groups

The variant allele frequency (VAFs) of SNVs (see methods) ranged from 0.23% to 96.47%, and, surprisingly, the majority of unique SNVs had a VAF below the current 1% detection limit (Fig. 1c & d). Functional annotation showed that 24.11% of SNVs were missense variants and high-frequency SNVs were enriched in the *CYTB* gene and D-loop regions (Fig. 1d). The mean SNV load (number of SNVs per genome, calculation in methods) among the individual mtDNA was 21.72 (Fig. 1e). The mutation spectra strongly biased toward transitions (i.e., A>G (T>C) and C>T (G>A)) (Supplementary Fig. 1c), which were consistent with previous reports^13, 17, 20^. The extensive-coverage UMI groups (covering > 95% of the full-length mtDNA with depth ≥ 3) supported de novo assembly of 11 full-length mtDNA. All of the 11 mtDNA were placed into the U5a1 mitochondrial haplogroup by MITOMASTER^21^, which fit the Dutch origin of 293 cells^22^, thus proving the accuracy of the assemblies.

Compared to Nanopore sequencing, PacBio circular consensus sequencing (CCS) generated 1.5X number of UMI groups from 2X number of reads, but the N50 was reduced by more than 2X and no UMI group covered > 95% of the full-length of mtDNA (Table 1, Fig. 1b). There were 637 high-confidence UMI groups containing 9,368 SNVs with frequencies from 0.16% to 100.00% (Fig. 1c). Unlike the Nanopore data that supported de novo assembly of full-length mitochondrial genomes, the PacBio data resulted in none. All of the PacBio high-frequency SNVs (VAF ≥ 0.6) existed in the Nanopore data, and all of them are common SNPs (dbSNP-151), thus cross-validating these called variants (Fig. 1d). Importantly, the mutational spectra and signatures were highly consistent between the PacBio and Nanopore data, and showed no evidence of artifactual mutations in homopolymer regions, which were purported to be more error-prone in Nanopore sequencing (Fig. 4b, Supplementary Fig. 1d). These data suggest the variants identified by Nanopore iMiGseq are of high confidence.

We evaluated the extent of false-positive variants due to polymerase replication error in the barcoding step (the main source of errors in UMI consensus sequencing^23^) and determined that this type of error introduced roughly 1 mutation in 1,600 labeled full-length mtDNA (see methods and shown previously^19^), thus representing a miniscule fraction of the SNVs (0.004% in the Nanopore iMiGseq). To further validate the ultra-rare variants detected by iMiGseq, we performed digital droplet PCR (ddPCR) assays for five SNVs representing a wide range of VAFs (0.24% to 41.18%) in the Nanopore iMiGseq data. The three rare SNVs with a VAF <1% (detected in one UMI group) were confirmed by ddPCR (Fig. 1f). The VAFs obtained by ddPCR were comparable to iMiGseq. The two high-frequency SNVs showed consistent VAFs in the iMiGseq by Nanopore and PacBio (m.4429G>A, 16.71% in Nanopore and 15.08% in PacBio; m.15259C>T, 41.18% in Nanopore and 39.12% in PacBio), and were further validated by ddPCR (m.4429G>A, 38.29%; m.15259C>T, 44.46%, Fig. 1f). It is worth noting that the difference in the VAF of m.4429G>A between iMiGseq and ddPCR is potentially due to the different population of cells in the experiments and heterogeneity of mtDNA in 293T cells.

Because Nanopore sequencing showed high concordance with PacBio CCS, which rivalled Illumina NGS in accuracy^24^ (Supplementary Fig. 2a), and PacBio failed to generate sufficiently long reads to cover the whole mtDNA, we concluded that Nanopore-based iMiGseq could produce highly accurate and complete mtDNA sequences. These results provided a quantitative analysis of the genetic heterogeneity of mitochondria in human cells beyond the analysis of individual SNVs to include the linkage between SNVs in the mitochondrial genome for the first time (Supplementary Fig. 1e).

### iMiGseq reveals genetic heterogeneity of mtDNA in single mouse oocytes

Heteroplasmic mutations have been documented in healthy human oocytes and primordial germ cells ^25, 26^. Experimental data and population genetics modeling suggest heteroplasmies arisen in mature oocytes strongly influence the inheritance of mtDNA mutations in the offspring^2, 3^. Yet, many key questions remain unanswered, such as whether the ostensibly somatic mutations found in old individuals originated from low-level heteroplasmic mutations in the oocyte or arose de novo. Also, many conclusions are based on imputation of data from bulk sequencing somatic cells rather than actual measurements in single oocytes. We hypothesized that iMiGseq could help address these questions by providing a quantitative understanding of the genetics of mtDNA in single oocytes.

We thus applied iMiGseq to single oocytes of the NOD-*scid* IL2Rgamma^null^ (NSG) and C57BL/6 (B6) mouse strains. Each oocyte was subjected to the same iMiGseq protocol and sequenced using one MinION flow cell. The three NSG oocytes generated 1.09, 0.96 and 1.99 million reads, which led to 4,078, 1,165 and 19,697 high-confidence UMI groups, respectively (Table 1, Supplementary Fig. 2b). VAULT analysis based on the NSG mouse reference genome (GenBank: CM004185.1) identified 182, 134 and 237 SNVs from oocytes NSG_1, NSG_2 and NSG_3, respectively. No large SV was detected. The results showed that the vast majority of the mtDNA (based on extensive-coverage UMI groups) in the three NSG oocytes contained no variant, which fit the expectation of a homogenous female germline resulted from extensive backcrosses to an inbred strain (Table 1, Fig. 2a). Most SNVs had a nominal heteroplasmy level (NHL, no. of UMI groups with SNV/no. of UMI groups with ≥ 3 coverage in the SNV position) less than 1%, which is incidentally the detection limit of most published studies^3, 12–14^ (Fig. 2b, Supplementary Fig. 3a). It is worth noting that the NHL of the rarest SNVs (e.g., appear in one UMI group) tends to be more affected by the sequencing depth and may not be accurate. Interestingly, the three oocytes shared a very small fraction of the SNVs despite being from the same female (Fig. 2c), and the shared SNVs tended to have high maximum occurrence times (no. of times a SNV is discovered among UMI groups). For example, the m.1442A>G, m.5421G>A, and m.14027C>A variants, shared by all three oocytes, had an occurrence time of 40, 30, and 66, respectively, as opposed to the median of 1 for all variants. These observations were consistent with an independent germline bottleneck in oocyte lineages^2^.

**Fig. 2.**
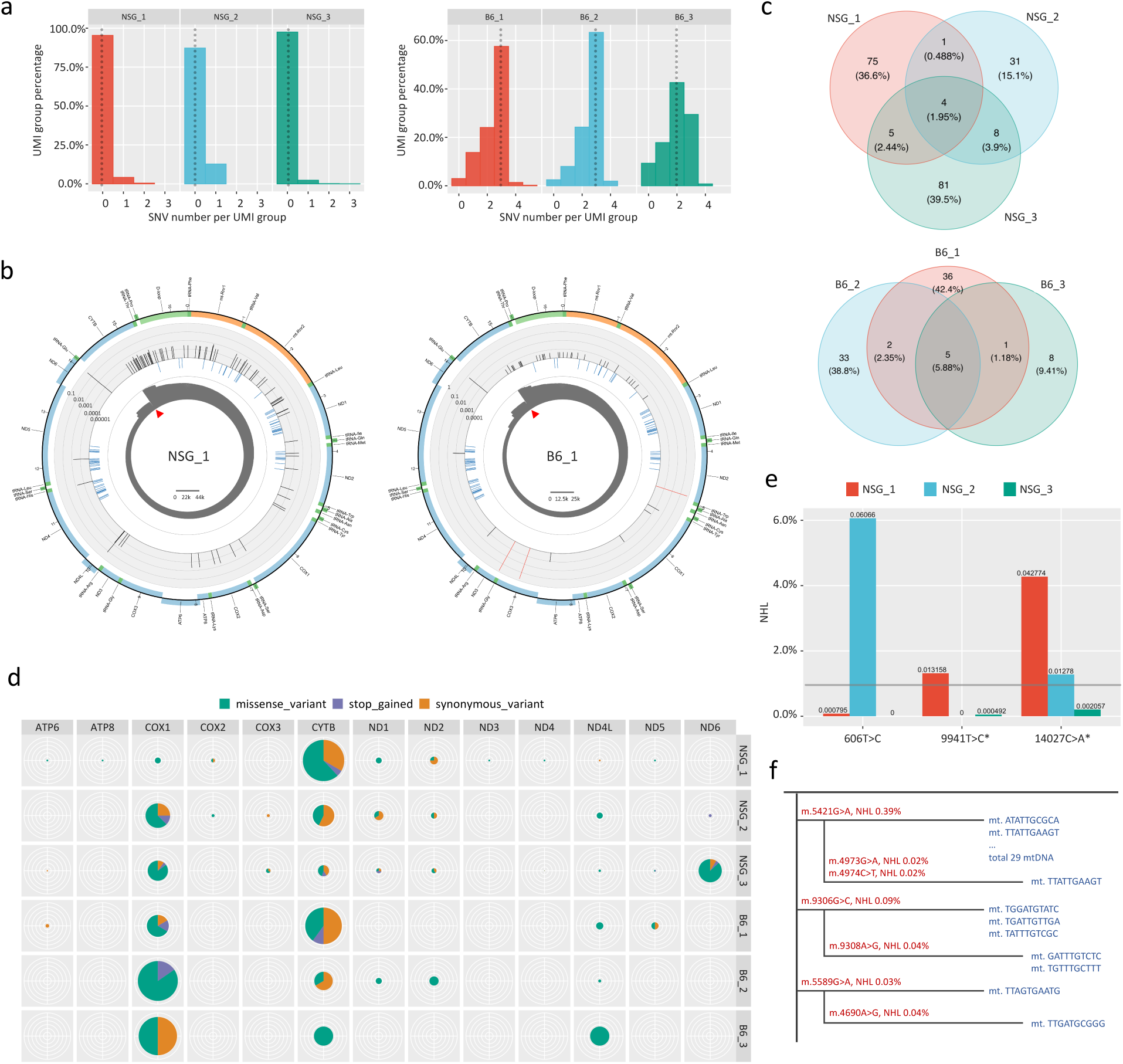
iMiGseq of single mouse oocytes. **a**, Histogram of SNV load per mtDNA in the three NSG (left) and B6 (right) mouse oocytes. The median SNV numbers are indicated by the vertical dotted line. The three oocytes of the same strain show a similar pattern of SNV load. **b**, Circular plots showing the distribution of SNVs in the mitochondrial genome of the three NSG (left) and B6 (right) mouse oocytes. The arrangement of the circular plots is similar to Fig. 1e, except that the middle circle (light blue) represents the common SNPs from the mouse dbSNP-142 database. **c**, Venn diagrams showing the overlap of mtDNA SNVs detected in the three NSG (upper) and B6 (lower) mouse oocytes. **d**, Functional annotation of SNVs in protein-coding regions detected by iMiGseq of single mouse oocytes. The size of the pie chart is proportional to overall mutation frequency, and the color corresponds to the type of mutation. Variants called in all UMI groups (not limited to extensive-coverage groups) were used to generate this plot. **e**, Comparison of common SNVs in the three NSG oocytes. * indicates detrimental variant. The detection limit of current NSG mtDNA sequencing is indicated by the horizontal line. **f**, Phylogenetic trees of individual detected mtDNA (shown in blue) constructed using ultra-rare SNVs (shown in Red) in the NSG_3 oocyte. The DNA sequences indicate the UMIs.

Functional annotation showed that deleterious SNVs (missense and stop-gained variants) affecting amino acid sequence were prevalent among heteroplasmic SNVs and accounted for 64.13 %, 17.17 %, and 51.06 % of SNVs discovered in NSG_1, NSG_2 and NSG_3, respectively (Fig. 2d, Supplementary Fig. 3b).

Similarly, we obtained individual mtDNA sequences and SNVs from three oocytes from one female of the B6 strain (Table 1, Fig. 2b, Supplementary Fig. 2c & 4a). Again, SV was not observed. As seen in the NSG data, only 5 (5.88%) SNVs were shared among the B6 oocytes (Fig. 2c). The majority of mtDNA in the three oocytes contained three high-frequency SNVs (NHL 42.80% to 72.82%, Fig. 2a), of which m.9027G>A caused a glycine to serine mutation in codon 141 of the *COX3* gene, while m.4891T>C and m.9461T>C were synonymous. The three SNVs were reported previously^27^. Like in NSG oocytes, deleterious SNVs in protein-coding sequences were prevalent among heteroplasmic SNVs in wild-type B6 oocytes (Fig. 2d, Supplementary Fig. 4b). Most low-frequency SNVs existed in solo UMI groups, however, several reoccurred in multiple UMI groups and/or oocytes, such as m.14027C>A (p.Gly15Val, in B6_1), m.10102G>A (p.Ala76Thr, in all oocytes), and m.5421G>A (p.Ala32Thr, in all oocytes).

The NSG and B6 reference mtDNA differ in the position m.9348 (B6/NSG:G/A), which served as the ground truth for estimating false discovery rate (FDR) of iMiGseq. Despite deep coverage (e.g., 10,876X) of UMI groups, this position showed zero non-reference SNV in either mouse strain, giving a conservative FDR estimate of < 9.2 × 10^-5^. We further took advantage of these variants to construct *in silico* ground-truth heteroplasmies by mixing various numbers of reads subsampled from NSG_2 with all reads of B6_2. iMiGseq accurately determined the heteroplasmy levels in iterated sub-samplings (71-105 times) even when the number of reads was limited (Supplementary Fig. 3e). These data were consistent with our previous data of IDMseq^19^ and showed that our methods provide accurate quantitation of the heteroplasmy levels that are reported in published mtDNA studies.

iMiGseq allowed us to compare the frequencies of variants shared by oocytes of the same mouse. We surveyed high frequency SNVs (> 1% NHL) in NSG oocytes and observed a significant difference in the frequency of the same variant (Fig. 2e). Two SNVs, m.606T>C and m.9941T>C (detrimental, p.Phe22Ser), showed ultra-low frequencies (< 0.1% NHL) in two oocytes but a drastically higher frequency in the third oocyte (Fig. 2e). These results suggested that ultra-rare detrimental variants could under some circumstances accumulate to frequencies above 1% during oogenesis, which supports the need for unbiased studies on these variants using ultra-sensitive methods such as iMiGseq.

The T>C (A>G) and G>A (C>T) transitions were predominant in all six mouse oocytes, consistent with known mtDNA-specific replication-coupled mutational signatures^13, 16, 28^ (Supplementary Fig. 3c, 4c & d). It is worth noting that the mutation spectrum of the SNVs with NHL< 1% was more diverse and could be influenced by one prevailing SNV (e.g., 14027C>A in NSG_1&3). The SNVs from extensive-coverage mtDNA were clustered in a heatmap to show haplotypes and linkage between heteroplasmic mutations (Supplementary Fig. 3d & 4e). Several phylogenetic trees were constructed based on haplotypes of individual mtDNA and showed evidence of sequential acquisition of ultra-rare SNVs (< 1% NHL), suggesting an accumulation of de novo mutations in individual mtDNA (Fig. 2f, Supplementary Fig. 3d). Together, the above findings revealed unappreciated levels of genetic heterogeneity of mtDNA in mouse oocytes and a surprisingly widespread presence of low-level deleterious variants due to mtDNA polymerase error. These haplotype-resolved variants supported an independent germline mtDNA bottleneck. These findings were only possible due to the analysis of individual mtDNA in single oocyte and would have been underestimated or missed by conventional methods.

### iMiGseq reveals genetic heterogeneity of mtDNA in single human oocytes

We next applied iMiGseq to three mature human oocytes (hOOs) from three donors. To achieve high UMI-labeling efficiency in single hOOs, we further screened for optimal primer sequences using mtDNA from five 293T cells (data not shown). iMiGseq captured 2,308, 1,461 and 1,535 UMI groups in hOO_1, hOO_2, and hOO_3, respectively (Table 1, Supplementary Fig. 2d). All three hOOs contained a significant amount of SNVs (75483 for hOO_1, 32276 for hOO_2, and 22522 for hOO_3). The NHL of SNVs separated into a high-frequency group (32.40%–97.18%) and a low-frequency group (0.04%–0.41%) (Fig. 3a, Supplementary Fig. 5a & b). The vast majority of SNVs had an NHL below the 1% detection limit of heteroplasmy reported in published studies^3, 12–14^ (Fig. 3b & c). These low-level heteroplasmic SNVs showed marked variations among the three donors, while the high-frequency SNVs (NHL > 1%) tended to highlight the shared genetic heritage of them (Fig. 3d). For example, the 11 SNVs shared by all three hOOs separate the L3 haplogroup from its ancestral L0 haplogroup. Interestingly, these SNVs are retained in Asian haplogroups M and F (a daughter group of N) but not in the European haplogroup H, which is consistent with the east Asian origin of the donors^21, 29^ (Supplementary Fig. 6). Note that haplogroup typing by iMiGseq is done in the context of full-length individual mtDNA in one assay as opposed to multiple genotyping assays as used currently.

**Fig. 3.**
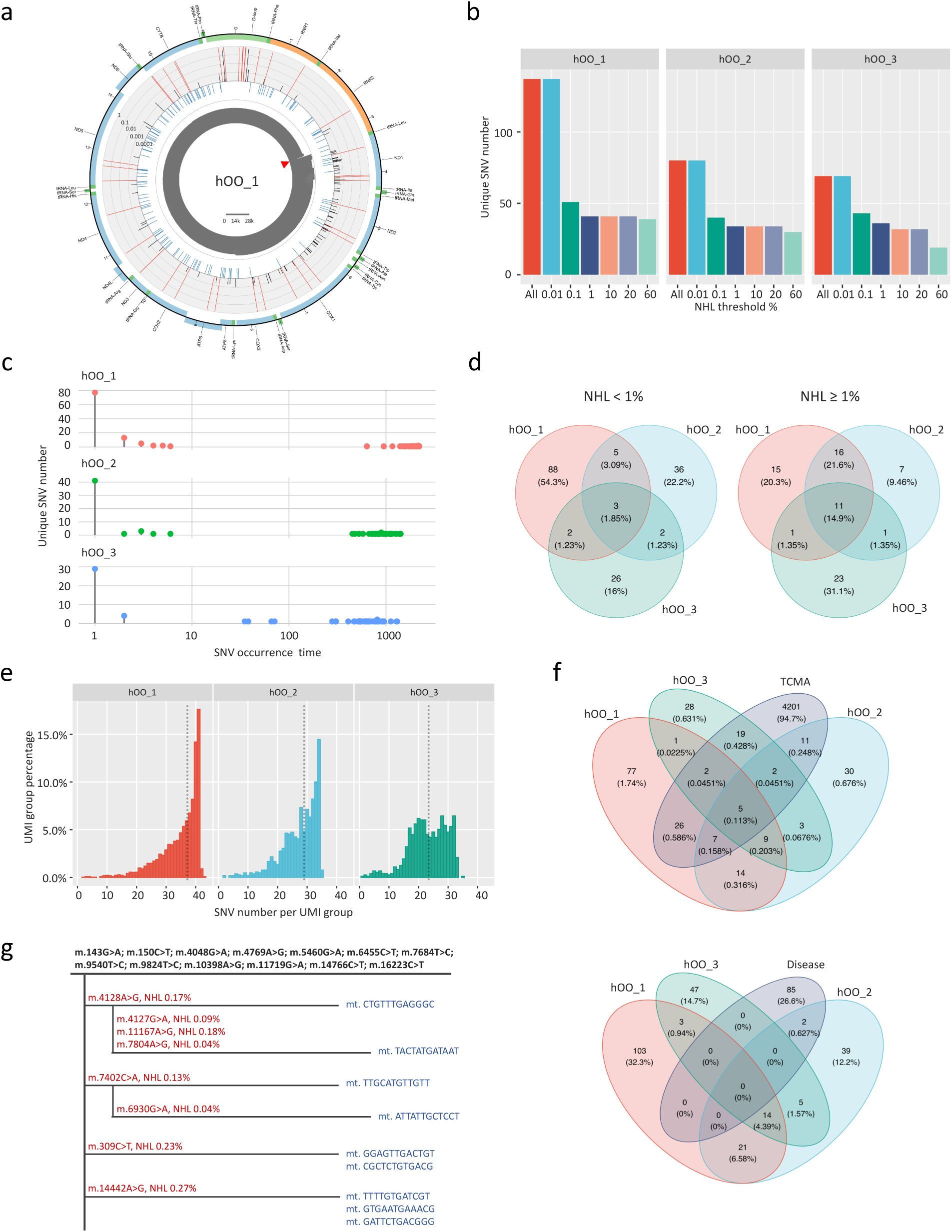
iMiGseq of single human oocytes. **a**, Circular plots showing the distribution of SNVs in the mitochondrial genome of hOO_1. The arrangement of the circular plots is similar to Fig. 1e. **b**, Unique SNV number above different heteroplasmy threshold. For all three hOOs, a significant amount of unique SNVs had an NHL below the 1% detection limit of current methods. **c**, Occurrence time of unique SNVs detected in iMiGseq of human oocytes. Most of the unique SNVs appear one time (exist in one UMI group). **d**, Venn diagrams showing the overlap of mtDNA SNVs between the hOOs at different heteroplasmy levels. A higher proportion of SNVs with an NHL ≥ 1% (right) were shared among hOOs than those with an NHL < 1% (left). **e**, Histogram of SNV load per mtDNA in the three hOOs. The median SNV numbers are indicated by the vertical dotted line. **f**, Venn diagrams showing the overlap of mtDNA SNVs in hOOs and cancers (TCMA, upper), or confirmed pathogenic mtDNA SNVs in mitochondrial diseases (Disease, lower). Cancer-related mtDNA SNVs exist in all three hOOs. **g**, Phylogenetic trees of individual detected mtDNA (shown in blue) constructed using ultra-rare SNVs (shown in Red) in the hOO_1 oocyte. Common SNVs (NHL > 1%) of the indicated mtDNA are shown in black. The DNA sequences indicate the UMIs.

We assembled consensus mtDNA sequences using reads from extensive-coverage UMI groups and obtained 125, 81 and 46 full-length mtDNA for hOO_1, hOO_2, and hOO_3, respectively (Supplementary File 1). MITOMASTER^21^ assigned all assembled mtDNA of hOO_2 and hOO_3 to the M7c1c2 and F1a1a haplogroups, respectively. For hOO_1, 122 assembled mtDNA were assigned to M7b1a1b, while the remaining 3 contained a different allele (m.3483G>C vs. A) consistent with the M7b1a1 haplogroup (Supplementary File 2). The m.3483G>C SNV caused the 59th amino acid to change from Glu to Asp in the *ND1* gene, and was also found in Genbank sequences (MF058561.1 and MF381578.1). Importantly, all of the assigned haplogroups matched the ethnic origin of the donors (Supplementary Fig. 6). These data show that iMiGseq provides base-resolution haplotypes of individual mtDNA in single cells.

The distribution of SNV load per mtDNA of healthy human oocytes was similar to that of 293T cells but distinct from mouse oocytes (compare Fig. 1e, 2a, and 3e). This is presumably due to the higher genetic diversity of mtDNA in outbred humans than in inbred mice. Previous reports based on deep NSG sequencing of human mtDNA identified several conserved mtDNA replication related regions (e.g., MT-LSP) that are devoid of even low-level heteroplasmy (1% detection limit)^7, 8^. Interestingly, iMiGseq identified two SNVs in such “variant-deserts” in two of four human samples with NHLs of 0.24% (m.438C>A in 293T cells) and 0.09% (m.16427C>T in hOO_2). Thus using the unbiased ultra-sensitive method iMiGseq, our data suggested that mtDNA mutations can randomly happen in any mtDNA region, and that the heteroplasmies in “variant deserts” are probably too rare to be captured by conventional NGS, which is consistent with the view that mutations in these regions impair mtDNA propagation^7, 8^.

The Cancer Mitochondrial Atlas (TCMA^13^) provided a comprehensive collection of heteroplasmic mutations in human cancers. However, the ontogeny of these cancer-associated mtDNA mutations whose occurrence positively correlates with age^13^ remains unclear. Interestingly, each of the three hOOs shared a significant portion (28.4%, 40.6% and 30.9% for hOO_1, hOO_2, and hOO_3, respectively) of its unique SNVs with the TCMA database^13^ (Fig. 3f). We refined the comparison by separating unique SNVs according to their NHL (≥ 1% or < 1%), which showed that a significant amount of shared SNVs existed well below the 1% detection limit of most published studies (Supplementary Fig. 5c & 7).

Intersecting the iMiGseq SNVs with a list of confirmed pathogenic mtDNA mutations^21^ identified two SNVs causing Leber’s hereditary optic neuropathy (LHON) and mitochondrial encephalopathy, lactic acidosis, and stroke-like episodes (MELAS, 3376G>A/p.Glu24Lys in *ND1*), and chronic progressive external ophthalmoplegia (CPEO, 5690A>G in tRNA^Asn^), respectively (Fig. 3f, Supplementary Fig. 5c). Each was in one mtDNA in hOO_2 with NHL of 0.07% (Supplementary Fig. 5d). We pooled data from all three hOOs *in silico* and showed that the same two mtDNA with LHON/MELAS and CPEO variants respectively were still correctly identified (Supplementary File 3), demonstrating the excellent sensitivity of iMiGseq for extremely low heteroplasmic mutations. These data provided evidence that low-level disease-associated heteroplasmic mutations commonly exist in single healthy human oocytes and they may contribute to late-onset cancers or inherited mitochondrial diseases through selection and/or genetic drift^2, 15, 30, 31^.

As in 293T cells and mouse oocytes, the predominant base substitutions in human oocytes were T>C (A>G) and C>T (G>A), well-known signatures for mitochondrial mutations^13, 16, 28^ (Fig. 4a, Supplementary Fig. 8b). The frequency of reciprocal mutations (e.g. T>C and A>G on the light strand) showed a strand bias, which has been observed by others and attributed to the difference in replication timing of the light and heavy strands^28, 32^. The spectra of all mutations were consistent with previous data^17^ (Fig. 4b). Some mutational signatures (e.g., in the T>C category) were shared by the three hOOs, but the diversity between samples was more evident than that between the Nanopore and PacBio data of 293T cells. The variations in mutational signatures were especially evident for SNVs with NHL < 1%, as also seen in mouse oocytes (Fig. 4b, Supplementary Fig. 4d). Lastly, data from the highly sensitive iMiGseq showed that oxidative damage (causing C>A/G>T transversions) is not a major source of de novo mtDNA mutations in the human germline. A similar observation was also made in somatic tissues^16^.

**Fig. 4.**
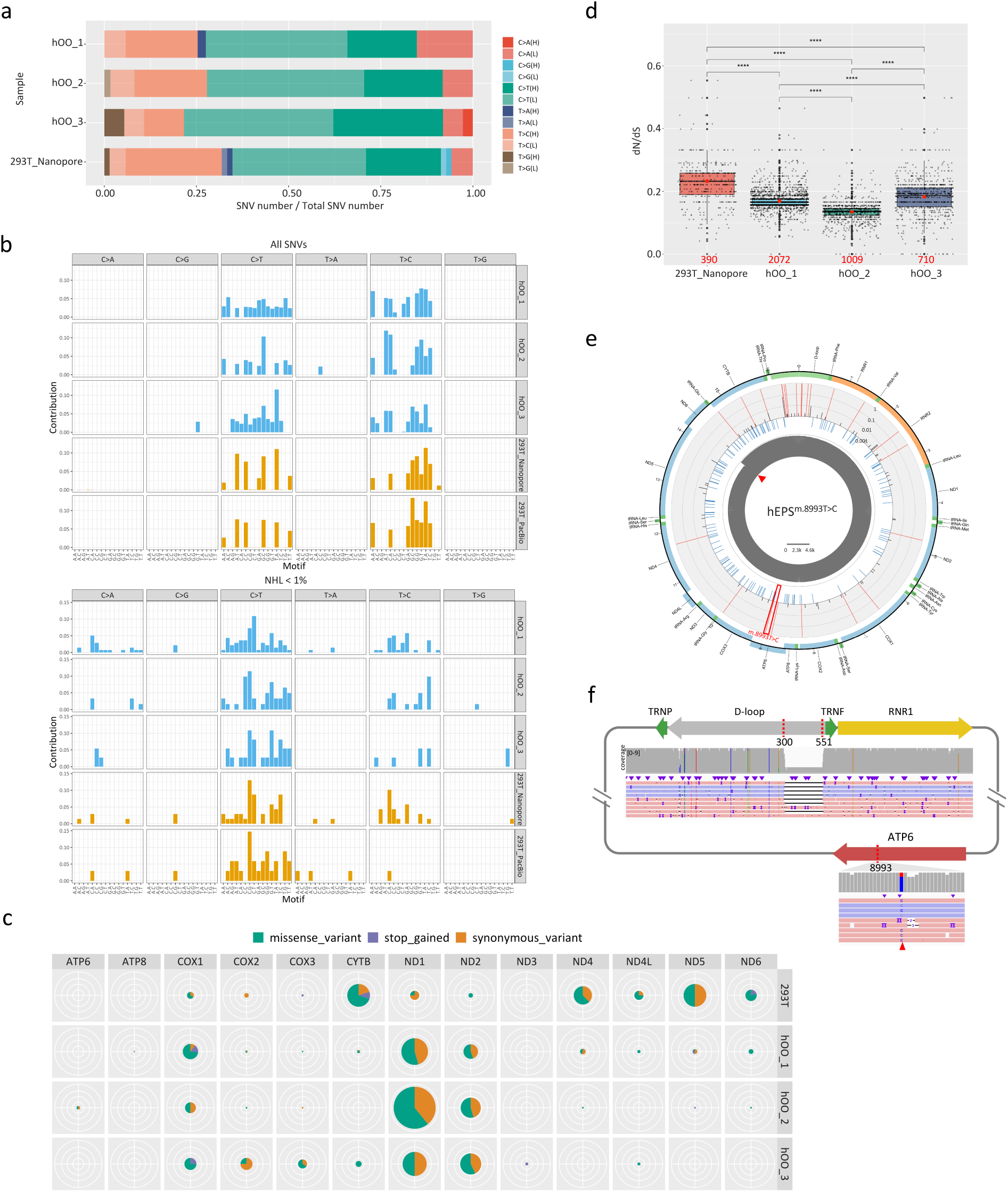
Analysis of mtDNA SNVs detected in human oocytes. **a**, Frequency of base substitutions on the heavy (H) and light (L) strands showing the strand asymmetry of low-frequency mtDNA SNVs (NHL < 1%) in hOOs and 293T cell line (sequenced by Nanopore). **b**, Highly consistent mtDNA mutational spectrum in hOOs and 293T cell line. The upper panel is for all SNVs, while the lower one is for SNVs with NHL < 1%. **c**, Functional annotation of SNVs in protein-coding regions detected by iMiGseq of hOOs and the 293T cell line (sequenced by Nanopore). Variants called in all UMI groups (not limited to extensive-coverage groups) were used to generate this plot. All hOOs show a similar enrichment of SNVs in *COX1*, *ND1* and *ND2* genes. The 293T cell line has more SNVs in the *CYTB*, *ND4* and *ND5* genes. **d**, Box plots showing the distribution of dN/dS ratio of individual mtDNA in hOOs and 293T cells (sequenced by Nanopore). The mtDNA numbers of every sample are shown (red values). The red dots inside the boxes indicate the mean values of dN/dS, and the thick black lines represent the median values. Potential outlier values are marked by bold black dots, while individual values calculated from every UMI group are shown as small grey dots. The lower and upper boundaries of the box represent the 25th and 75th percentiles, respectively. The p values were based on the Kruskal-Wallis test, **** stands for p< 2.22e-16. **e**, Circular plots showing the distribution of SNVs in the mitochondrial genome of hEPS^m.8993T>C^. The arrangement of the circular plot is similar to Fig. 1e and the m.8993T>C mutation is indicated by the red frame. The data were generated from thirty hEPS cells. **f**, A schematic map of the mutant mtDNA harboring the 251-bp deletion and m.8993T>C mutation detected in hEPS^m.8993T>C^ and the accompanying Integrative Genomics Viewer (IGV) tracks showing the alignment of Nanopore reads.

No enrichment or depletion of low-NHL SNVs (< 1%) was observed in the D-loop and tRNA genes in any of the hOOs (Supplementary Table 1). Mutations in rRNA genes were significantly depleted in two of the three hOOs (the samples number of hOO_3 was too small for proper statistical testing, Supplementary Table 1). Among protein-coding genes, the three hOOs consistently showed frequent mutations in the *ND1*, *ND2,* and *COX1* genes, which was a different pattern compared to cultured 293T cells (Fig. 4c). Deleterious SNVs accounted for 24.10%, 21.47% and 21.54% of the SNVs in hOO_1, hOO_2, and hOO_3, respectively (Supplementary Fig. 8a). Synonymous mutations occurred significantly more frequently than expected by chance, while non-synonymous mutations did not deviate from random chance (Supplementary Table 1).

The SNVs from extensive-coverage mtDNA were clustered in a heatmap (Supplementary Fig. 8c). As seen in mouse oocytes, phylogenetic trees constructed based on individual mtDNA haplotypes showed evidence of sequential acquisition of ultra-rare SNVs (< 1% NHL), suggesting an accumulation of de novo mutations in individual mtDNA (Fig. 3g). The dN/dS ratio is a major tool used to detect selection on protein-coding mutations^33^, and has been used to study selection on mtDNA^13, 28, 34^. The dN/dS ratios of all mtDNA in each of the three hOOs showed strong evidence for purifying (negative) selection against non-synonymous mutations (Fig. 4d). Considering that oocytes experience an mtDNA bottleneck during germ cell development^35^, our data support the role of the germline bottleneck in eliminating deleterious mutations in the offspring.

### iMiGseq quantitates mtDNA mutations in NARP/Leigh syndrome hEPS cells

Since iMiGseq enabled accurate quantitation of mtDNA mutations in a haplotype-resolved manner, we hypothesized that it could improve genetic diagnosis of mitochondrial diseases. Mouse extended pluripotent stem (mEPS) cells can generate blastocyst-like structures (EPS-blastoids) through lineage segregation and self-organization *in vitro*^36^. If an analogous system can be established using human EPS (hEPS) cells, it could provide a unique model of mitochondrial disease during early embryogenesis. Therefore, as a proof-of-concept, we performed iMiGseq in hEPS cells derived from mitochondrial disease-specific induced pluripotent stem cells (iPSCs) described previously^37^. The patient is a carrier of the mtDNA mutation (m.8993T>C) for neuropathy, ataxia and retinitis pigmentosa (NARP) and maternally inherited Leigh syndrome. She gave birth to two daughters who died of Leigh syndrome and one son who was age three and asymptomatic at the time of sample collection, all confirmed carriers of m.8993T>C. The NAPR/Leigh syndrome hEPS cells (referred to as hEPS^m.8993T>C^ hereafter) were collected for UMI labeling of whole mtDNA with different primer designs (important for avoiding primers overlapping with disease relevant SNPs) (Supplementary Table 2). As in hOOs, iMiGseq showed an extensive coverage of full-length mtDNA in the hEPS^m.8993T>C^ that supported detailed analysis of variants (Fig. 4e, Supplementary Fig. 9a-c, and Table 1). It detected the m.8993T>C mutation in 74.2% of mtDNA, which matched the heteroplasmy levels observed in the patient’s oocytes (70∼90%, Fig. 4e). Interestingly, one of the m.8993T>C mutant mtDNA also harbored a 251-bp deletion in the D-loop region (Fig. 4f). These results clearly demonstrated that iMiGseq could not only detect and quantitate disease-causing mutations but also provide comprehensive characterizations of all types of mutations, including SNVs and SVs, and their genetic linkage with high sensitivity and specificity.

## Discussion

To the best of our knowledge, these data represent the first demonstration of an unbiased high-throughput base-resolution analysis of individual full-length mtDNA in single cells. Taking advantages of molecular consensus sequencing (IDMseq) and a specially designed bioinformatics pipeline (VAULT), iMiGseq greatly improved the sensitivity of heteroplasmy detection and showed that most unique mtDNA SNVs in cells are rare and well below the current 1% detection limit. They suggest that a hitherto underestimated population of rare de novo mtDNA variants exist in the female germline, much like the submerged portion of an iceberg hidden below the technological limit. The analysis of oocytes from the same female mouse showed ultra-rare detrimental variants can accumulate and increase frequencies to above 1% during oogenesis. During oogenesis mtDNA experiences a genetic bottleneck that could result in large swings in heteroplasmy levels between mother and child or between offspring^2, 3, 38^. It is conceivable that some of the rare detrimental variants can resurface due to genetic drift, mtDNA bottleneck, selection or a combination of any of these processes and become high-level heteroplasmic variants in human pedigrees and to the development of complex diseases during aging^4, 39^.

Further analysis showed that deleterious SNVs were prevalent among heteroplasmic SNVs in both mouse and human oocytes. A sizable portion of low-level heteroplasmic SNVs are associated with cancers or mitochondrial diseases, and the phylogenetic trees of individual mtDNA suggest a sequential accumulation of de novo mutations (that are rare at the population level). These observations raise important questions about the roles of these rare mtDNA mutations that exist at the beginning of one’s life, and their implications for the diagnosis and prevention of mitochondrial diseases. Note that we only surveyed one of the most conservative lists of mitochondrial disease variants and many other mtDNA variants have been reported to link to mitochondrial diseases and complex diseases^4, 21^, so the overlap with disease variants may be even bigger. iMiGseq could offer new opportunities to follow the dynamics of such SNVs during development and aging in hope of deciphering the emergence of mutations and understanding their clinical significance.

Because NGS of mtDNA necessitates fragmentation of mtDNA molecules, it is impossible to ascertain the true haplotype of individual mtDNA molecules. It is thus extremely difficult to use short-read data to study genetic interaction between rare mutations and their genetic backgrounds. Short reads also could be erroneously mapped to NUMTs, causing false variants^11^. Taking advantages of ultra-long reads of Nanopore sequencing, iMiGseq could provide thousands of full-length mtDNA and their variants in a cell, which completely avoids NUMTs and enables studies of interactions between different heteroplasmies. Nanopore-based IDMseq has been shown to offer superior characterization of SVs induced by CRISPR/Cas9 editing in human embryonic stem cells^19^. Interestingly, unlike widespread low-level heteroplasmies uncovered by iMiGseq, no large SV was detected in healthy cells. One limitation of the current analysis is variant calling for small insertions and deletions, which remains challenging in most sequencing platforms^40, 41^. Although the sample sizes were limited, our high-sensitivity data provide strong evidence that SVs are rarely transmitted through the human germline^4^, and suggest a strong purifying selection against deleterious SVs in oogenesis.

Because iMiGseq enables for the first time quantitative base-resolution analysis of thousands of mtDNA in single cells, it offers an unprecedented opportunity for preimplantation genetic diagnosis (PGD) of mitochondrial diseases. Despite the prevalence (∼1 in 5000) of mitochondrial diseases, current diagnostic procedures have been ineffective due to large genetic heterogeneity of mtDNA among different cells. iMiGseq could be used to analyze mtDNA mutational load in single biopsied blastomeres, which have been shown by several studies^42–45^ to faithfully represent the heteroplasmy level of in vitro fertilized embryos. Our proof-of-concept data in hEPS^m.8993T>C^ show that this is not only feasible but also likely to yield novel knowledge of the full spectrum of mutations and their genetic linkage. Besides the germline, it is logical to extend iMiGseq technology to somatic tissues to unravel the direction of causality between mtDNA mutations and aging and complex diseases in the future. Similarly, because iMiGseq works for different species and cell types, we expect it to be widely applicable to many fields for the study of this ancient organelle that energizes most life forms on earth.

## Materials and Methods

### Cell lines and human oocytes

The 293T cell line was purchased from ATCC and cultured in Gibco™ DMEM medium (high glucose) containing 10% Gibco™ Fetal Bovine Serum (heat inactivated) and 1X Gibco™ Penicillin-Streptomycin (5,000 U/mL). Cells were maintained at 37°C in a humidified incubator with sea-level air enriched with 5% CO_2_. All human immature oocytes were collected from the reproductive medical center in the Third Affiliated Hospital of Guangzhou Medical University. The study of human oocytes collection was approved by the Institutional Review Board (IRB) of the Third Affiliated Hospital of Guangzhou Medical University and KAUST Institutional Biosafety and Bioethics Committee (IBEC). All of the oocyte donors signed the inform consent voluntarily after they were clearly informed all of the content and details of the experiments. The women were in intracytoplasmic sperm injection (ICSI) cycles because of male infertility were involved in this study, and the immature oocytes were found after oocyte retrieval and cumulus cells removal. In general, such immature oocytes will be discarded as a medical waste and not used for the fertilization during the process of assisted reproductive technology. The endometrium tissues were donated by women with mitochondria diseases who signed the informed written consent voluntarily after learning the study aims and methods. The protocol was approved by the Institutional Review Board of Peking University Third Hospital. Endometrium tissues were collected into a 50 ml centrifuge tube with DMEM/F12 culture media (no phenol red, Invitrogen, Carlsbad, CA, USA) containing 100 U/ml penicillin and 100 μg/ml streptomycin (Invitrogen) during a routine gynecological examination. The tube with endometrial sample was put in the ice, and quickly transported to the laboratory. The endometrial tissue was rinsed using Hanks buffered salt solution (HBSS, Invitrogen) three times and minced into 1 mm^3^ fragments. After digested with 2 mg/ml collagenase type I (Life Technologies, New York, NY, USA) for 45 minutes and DNase I for 30 min subsequently, the dissociated cellular suspension was treated with ACK lysis buffer (Life Technologies, New York, NY, USA) to remove red blood cells. The cells were then centrifuged for 200 g for 10 minutes, and resuspended using DMEM/F12 supplemented with 10% fetal bovine serum (Charcoal/Dextran Stripped, Gemini, California, USA), and plated on 35 mm dishes (Corning) at 37 °C in 5% CO_2_.

### In vitro maturation of human oocytes

Human immature oocytes were cultured following our previous protocols which was consist of IVM basal medium, 0.075 IU/ml FSH, 0.075 IU/ml luteinizing hormone (LH), 10 ng/ml EGF, 10 ng/ml BDNF and 10 ng/ml IGF-1^46^. The oocytes expelling the first polar body were regarded as mature oocytes at metaphase II stage. Single matured MII oocyte was transferred into 0.5ml EP tube without medium using mouth pipette and frozen in −80°C refrigerator before further molecular analysis.

### Mouse oocyte isolation

The animal experiments in this study were approved by the Institutional Animal Care and Use Committee (IACUC) of KAUST and the Salk institute for biological studies. The NSG and C57BL/6 mice were purchased from the Jackson Laboratory and Charles River Laboratories and kept in KAUST animal resources core lab. The NSG oocytes were collected from naturally ovulating female mice. For B6 oocytes, superovulation was induced in C57BL/6 (6 weeks) female mice by sequential intraperitoneal injection of 5 international units (IU) of pregnant mare’s serum gonadotrophin (PMSG) (USBiological Life Sciences G8575A) and 5 IU of human chorionic gonadotrophin (hCG) (Sigma-Aldrich C1063) 46-48 h later. C57BL/6 mice were sacrificed 14 h after hCG injection. Oviducts were dissected from NSG mice (16-20 weeks) or from the super-ovulated C57BL/6 mice, and the oocytes were isolated by mouth pipette and washed in cold PBS. Then, the oocytes were dissociated from cumulus cells using Accutase (5-10 min RT), washed in cold PBS by pipetting up and down, transferred to PCR tubes in a small volume of PBS, and frozen at −80°C.

### Derivation and culture of hEPS cells

To generate mitochondria diseases iPSCs, endometrium tissues fibroblast cells were transfected with a Sendai virus reprogramming kit (Life Technologies, A16517). The transfected cells were then plated onto Matrigel-coated culture dishes according to the manufacturer’s instructions. The iPSCs were cultured on Matrigel-coated tissue culture dishes (ES-qualified, BD Biosciences) with mTeSR1 (STEMCELL Technologies) at 37°C and 5% CO2 in a humidified atmosphere incubator. The iPSCs culture medium was changed daily. The cells were passaged every 3–4 days using Accutase (Stemcell Technologies). The iPS cells conversion to EPS as previous reported^47^.

### ddPCR

ddPCR was performed on a Bio-Rad QX200 Droplet Digital PCR System using ddPCR Supermix for Probes (No dUTP) kit (Bio-Rad, 1863024) according to manufacturer’s protocols. The probes were synthesized by Integrated DNA Technologies Inc. as PrimeTime qPCR Probes. The wildtype probes were labeled as 5’HEX/ZEN/3’IBFQ, while the mutant probes were labeled by 5’FAM/ZEN/3’IBFQ. The primer and probe sequences were shown in Supplementary Table 2. The primer/probe ratio was set as 3.6:1. For each reaction, 0.5 ng of purified 293T cell genome were used. All experiments were performed in three independent replicates, and the positive events were combined for the analysis.

### Cell lysis and DNA purification

Five 293T cells, single oocytes, or 5 to 30 previously frozen hEPS^m.8993T>C^ cells were pelleted by centrifuge at 200g for 3 mins, and then lysed in 5 µl RIPA buffer (150 mM sodium chloride, 1.0% NP-40, 0.5% sodium deoxycholate, 0.1% sodium dodecyl sulfate, 50 mM Tris, pH 8.0) on ice for 15 mins. The cell lysate was diluted by adding 10 µl H_2_0, and further purified to extract total DNA with 15 µl Beckman Coulter AMPure XP beads (A63882). Two-round of 70% ethanol washes were performed to remove detergents. The DNA was eluted in 10 µl H_2_O to be used for the UMI labeling of mtDNA.

### UMI labeling and amplification of mtDNA

The targeted UMI labeling of individual mtDNA was achieved by mtDNA specific UMI oligos. The oligos were selected to enable the efficient amplification of full-length human or mouse mitochondrial genomes. A 5’ universal primer sequence and middle UMI sequence were added to the five-prime end of mtDNA specific oligos to form UMI oligos. The full list of oligos used in this study is shown in Supplementary Table 2. The BLAST of human primers to the human reference genome showed that no primers will amplify NUMTs. Since iMiGseq amplified the whole circular mtDNA genome using inverse primers, linear NUMTs were less likely to be amplified by our inverse primers. The data analysis can easily detect and exclude NUMTs since they share a different structure than the full-length mtDNA. A screen of DNA polymerase was performed to ensure the high efficiency in the UMI labeling and PCR amplification.

The UMI labeling reaction was set up as follows: 10 µl purified DNA, 2.5 µl UMI oligos (10 µM), 12.5 µl 2X Platinum™ SuperFi™ PCR Master Mix (Invitrogen, 12358010). The reaction was incubated on a thermocycler with a ramp rate of 1°C per second using the following program: 98 °C 1 min, 70 °C 5 s, 69 °C 5 s, 68 °C 5 s, 67 °C 5 s, 66 °C 5 s, 65 °C 5 s, 72 °C 10 min, 4 °C hold. After UMI labeling DNA is purified by 0.8X AMPure XP beads, washed twice by 70% ethanol and eluted 10 µl H_2_O. The universal primer and mtDNA specific reverse primer were used to amplify only the UMI labeled mtDNA. The 50 µl PCR reaction contains 1.25 U PrimeSTAR GXL DNA Polymerase (Takara, R050), 1X PrimeSTAR GXL Buffer, 200 µM dNTP mixture, 0.2 µM each primer, and 10 µl of purified UMI-labeled DNA. The thermocycler program is set as follows: 95 °C 1 min, (98 °C 10 s, 68 °C 14 min, 30 cycles), 68 °C 5 min, 4 °C hold. The amplicon was validated by agarose gel electrophoresis. If a specific DNA band was observed, the rest DNA would be purified by 0.8X AMPure XP beads. A second round of PCR amplification for 15-20 cycles with PrimeSTAR GXL DNA Polymerase could be performed to obtain enough DNA for sequencing.

We considered if the variant calling results could be affected by PCR artifacts, which is the main source of errors in UMI consensus sequencing originating from polymerase replication error in the barcoding step^23^. The Platinum SuperFi DNA polymerase we used has the highest reported fidelity (> 300X that of Taq polymerase), and meanwhile captures twice more molecules in the library than Taq^19^. Theoretically, this polymerase introduces one error in ∼1,600 unique 16556-bp molecules in the UMI labeling step. Accordingly, this type of inescapable error is expected to be around 1 in 1,600 UMI groups, thus representing a minor fraction of the observed SNVs.

### Library preparation and sequencing

For Nanopore sequencing, library preparations were done using the ligation sequencing kit (Oxford Nanopore Technologies, SQK-LSK109). The sequencing runs were performed on an Oxford Nanopore MinION sequencer using R9.4.1 flow cells. Base calling of Nanopore reads was done using the official basecaller termed Guppy (v3.2.1). For PacBio sequencing, library preparations were done using the Sequel Sequencing Kit 3.0. The sequencing runs were performed by the BIOPIC core facility at Peking University (Beijing, China) on the PacBio Sequel with SMRT Cell 1M v3 LR Tray. HiFi Reads were generated by the official tool termed ccs (v3.4.1). All procedures were preformed according to manufacturer’s protocols.

### Bioinformatic analysis

All iMiGseq data were analyzed by VAULT with *--unmapped_reads* and *--group_filter* options, to remove unmapped reads before UMI analysis and filter out low-confidence UMI groups after variants calling. The percentage of reads with UMI depends on base calling errors in the UMI region and/or DNA fragmentation during library preparation and sequencing. For Nanopore data, the error tolerance threshold *--error* in UMI identification was set to 0.11, while for PacBio CCS data, it was set to 0.05. Only perfect UMIs with correct length and structure (NNNNNTGNNNNN) were subjected to downstream analysis. The SNV calling and filter were performed using default parameters of VAULT, and involved in Samtools v1.9^48^. SNVs in primer regions were filtered out before downstream analysis.

The reference sequence *--refer* used in VAULT analysis was the designed amplicon sequence, which is based on CM004185.1 for NOD oocytes, NC_005089 for B6 oocytes, and NC_012920 for 293T cells and human oocytes. This gave rise to a different coordinate to the canonical mitochondria reference genome. The *vault position* command was used to revise the DNA coordinate and reference chromosome name in VCF files, to enable further functional analysis by SnpEff v4.3^49^. In the SNV annotation, the positions of SNVs in NOD oocytes were converted to the corresponding coordinates of B6 mouse strain. GRCm38.86 database of SnpEff was used in mouse SNV annotation, while hg38kg database was used for human SNVs.

The SV calling of VAULT utilized minimap2.1^50^ and sniffles v1.0.11^51^. The detected SVs were first filtered by a variant allele frequency of 0.6 and then manually checked. This SV calling pipeline had been validated in the original work of VAULT^19^. However, in this study we didn’t detect any large SVs (≥35 bp) from mtDNA of healthy samples. The consensus sequence of high coverage UMI groups was called using *vault consensus* command. It utilized canu v2.0^52^ to do de novo assembly, and the Nanopore official tool medata v0.12.1 to polish the assembled sequence. Most UMI groups failed to generate contigs in canu assembly, thus lead to a reduction of assembled mtDNA. Further assemblers with improved performance could potentially solve this problem and lead to more assembled mtDNA.

The synonymous and nonsynonymous sites of mouse mitochondrial genome was calculated using the *pS_pN_count* tool of VAULT, and based on the Nei-Gojobori method^53^. The dN/dS ratio was calculated using bash command based on the Nei-Gojobori method. The mutational spectrum was analyzed using SomaticSignatures^54^.

SNV allele frequency (VAF, also mentioned as NHL) is calculated as

*UMI group number with this SNV / effective UMI group number at this position*

Effective UMI group number is defined as the number of UMI groups with depth ≥ 3 at the position of this SNV.

SNV number per genome is calculated as

*SNV number in UMI group / surveyed length (depth ≥ 3) in that group * genome length*.

## Data and materials availability

VAULT and sample data in this study are accessible at GitHub (https://github.com/milesjor/vault). Raw sequencing data and oligo sequences are available upon reasonable request.

## ACKNOWLEDGEMENTS

We thank members of the Li laboratory, Khaled Alsayegh, Samhan Alsolami for helpful discussions; Jinna Xu and Marie Krenz Y. Sicat for administrative support. We thank Chenyang Geng at the BIOPIC core facility at Peking university for technical assistance in PacBio sequencing. We thank Professor Jasmeen Merzaban’s lab and KAUST Animal Research Core Laboratory for sharing mouse strains. The research of the Li laboratory was supported by KAUST Office of Sponsored Research (OSR), under award number BAS/1/1080-01. The work was supported by a KAUST Competitive Research Grant (award number URF/1/3412-01-01) given to ML, YH and JCIB; the National Key R&D Program of China (2016YFC1000601) and the National Natural Science Funds (81571400, 81771580 and 81971381), MMAAP foundation to YY; and the National Key Research and Development Program of China (2019YFA0110804, 2018YFC1003203), and the National Natural Science Foundation of China (81871162) to YF.

## Author contributions

CB and LW performed majority of the experiments related to sequencing. YF performed the human oocyte collection and experiments related to hEPS cells. CB performed the bioinformatics analysis. CB and JW performed experiments related to PacBio sequencing. CB, JCIB, and ML analyzed the data and wrote the manuscript. YH and YY contributed to the writing of the manuscript. CB and ML conceived the study. YY, JCIB and ML supervised the study.

## Competing interests

A patent application based on methods described in this paper has been filed by King Abdullah University of Science and Technology, in which CB, LW and ML are listed as inventors. The authors declare no other competing interest.

## Supplemental figures

**Supplementary Fig. 1.**
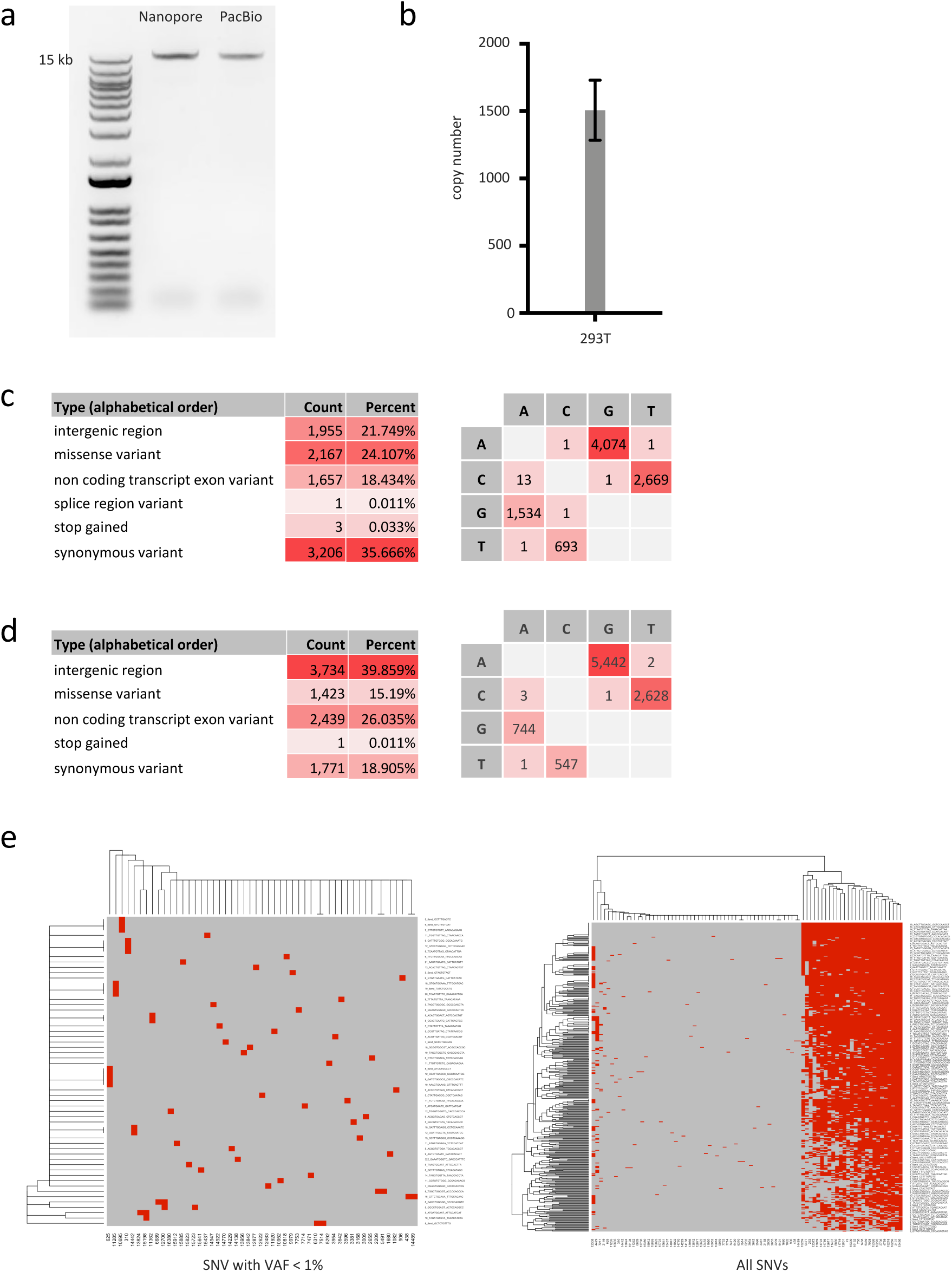
iMiGseq for five 293T cells. **a**, Agarose gel picture showing full-length mtDNA amplified in iMiGseq of 293T cells. **b**, qPCR quantitation of mtDNA copy number of 293T cells. The average mtDNA copy number is 1507 per cell. **c**, Analysis of SNVs detected by Nanopore iMiGseq based on functional annotation and base change. The majority of base changes are A>G (T>C) and C>T (G>A). **d**, Similar to **c** but for PacBio iMiGseq data. **e**, Heatmaps of clustering of individual mtDNA detected in 293T cells by Nanopore iMiGseq based on their genotype. The analysis of low VAF SNVs (NHL (nominal heteroplasmy level, see main text) < 1%) is shown on the left, and that of all SNV is on the right.

**Supplementary Fig. 2.**
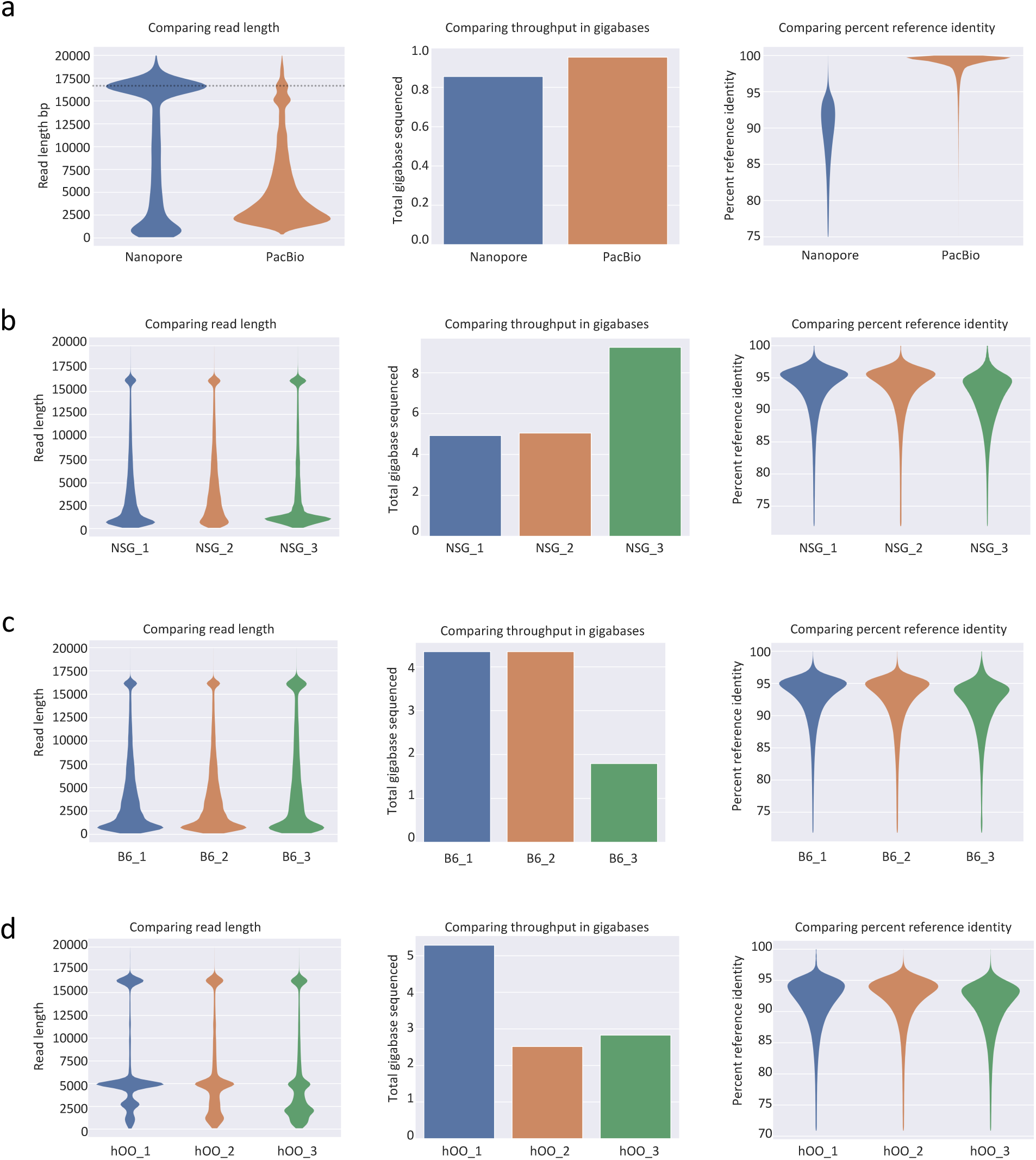
Quality control of reads in all sequencing runs in this study. **a**, Comparison of read length, throughput and percent reference identity of reads in the 293T sequencing by Nanopore and PacBio sequencers. PacBio ccs reads have a higher accuracy than Nanopore 1D reads. **b**, Comparison of read length, throughput and percent reference identity of reads in the NSG mouse oocyte sequencing. **c** and **d**, Similar to **b** but for the B6 mouse oocyte and human oocyte sequencing, respectively.

**Supplementary Fig. 3.**
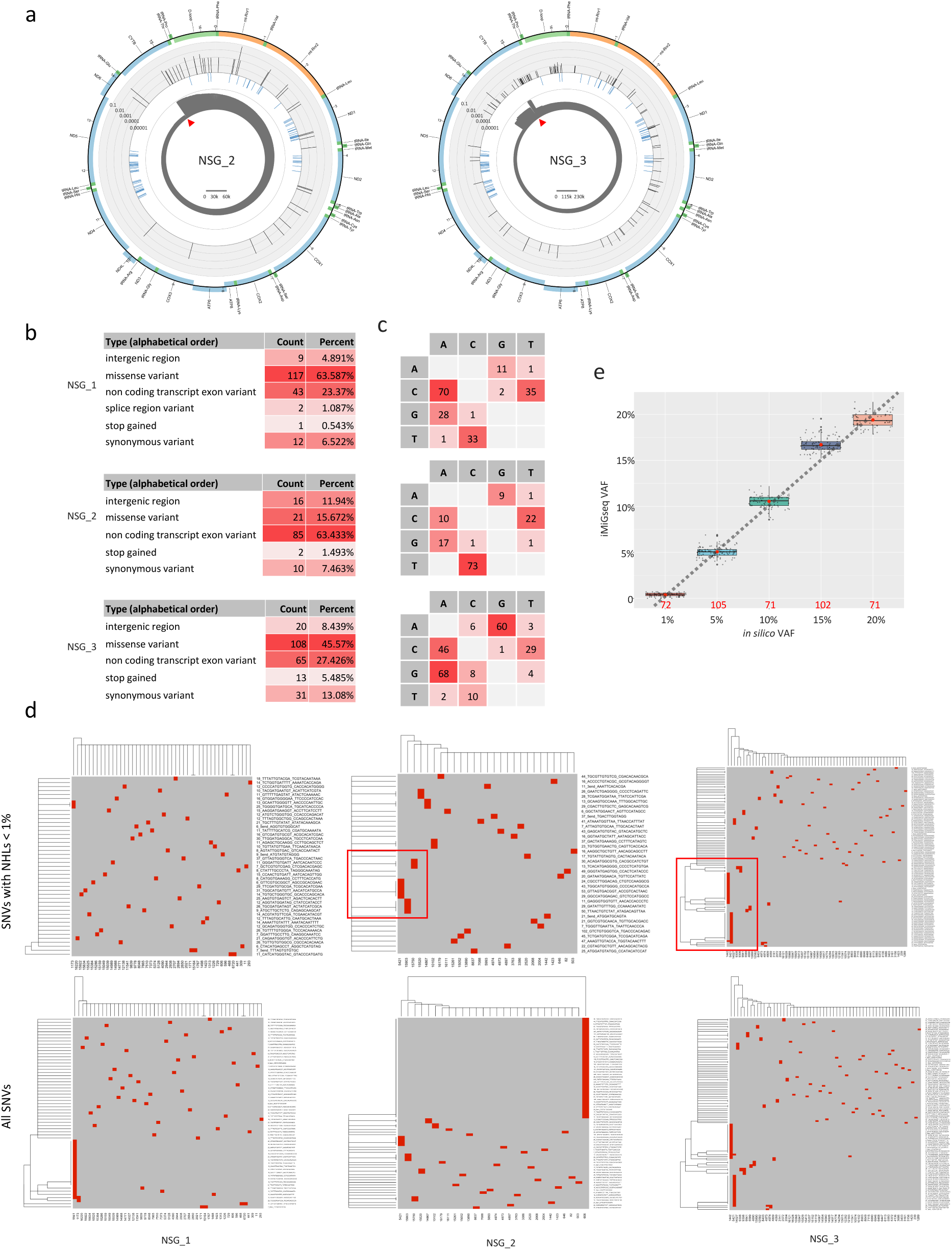
SNV analysis in the iMiGseq of NSG mouse oocytes. **a**, Circular plots showing the distribution of SNVs in the mitochondrial genome of NSG_2 and NSG_3 oocytes. The arrangement of the circular plots is similar to Fig. 2b. Please note that the NSG_3 sequencing experiment was carried out with an 8N UMI instead of 10N, which resulted in higher recovery of UMI groups. **b**, Analysis of mtDNA SNVs of NSG mouse oocytes based on functional annotation. **c**, Analysis of mtDNA SNVs of NSG mouse oocytes based on base change. The majority of base changes are T>C (A>G) and G>A (C>T). **d**, Heatmaps of clustering of individual mtDNA detected in NSG mouse oocytes based on SNVs (top panels: NHL < 1%, bottom panels: all SNVs). The red boxes highlight several lineages of individual mtDNA with ultra-rare SNVs (< 1% NHL), some of which acquired additional de novo variants. **e**, Comparison between *in silico* VAFs and VAFs as determined by iMiGseq in different subsampling experiments. The box plots show the distribution of VAFs determined by iMiGseq for the respective *in silico* VAFs. The percentages are those of the NSG allele. The numbers of sub-sampling iterations are shown as red numbers. The red dots inside the boxes indicate the mean values, and the thick black lines represent the median values. Potential outlier values are marked by bold black dots, while individual values of sub-samplings are shown as small grey dots. The lower and upper boundaries of the box represent the 25th and 75th percentiles, respectively.

**Supplementary Fig. 4.**
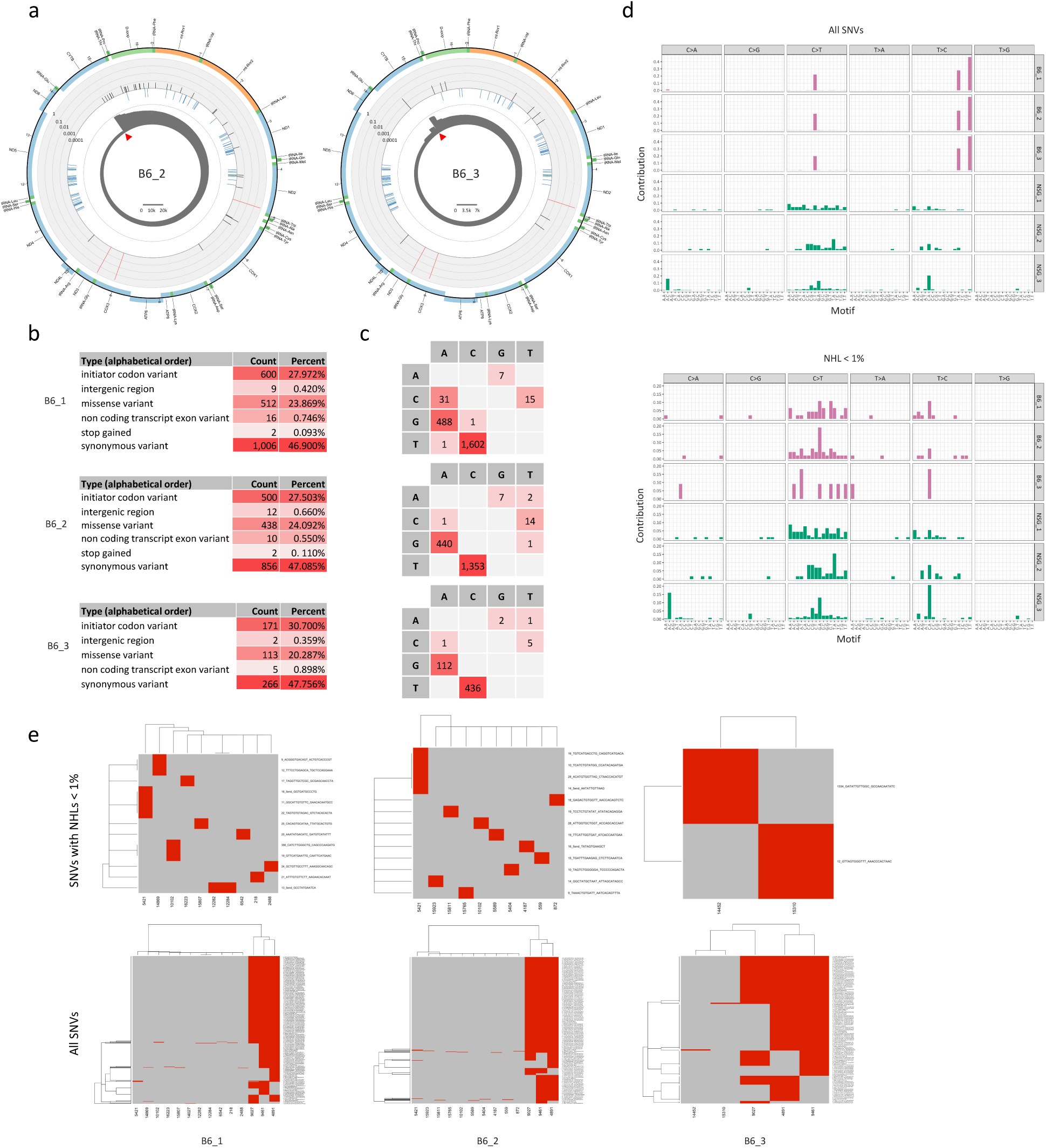
SNV analysis in the iMiGseq of B6 mouse oocytes. **a**, Circular plots showing the distribution of SNVs in the mitochondrial genome of B6_2 and B6_3 oocytes. The arrangement of the circular plots is similar to Fig. 2b. **b**, Analysis of mtDNA SNVs of B6 mouse oocytes based on functional annotation. **c**, Analysis of mtDNA SNVs of B6 mouse oocytes based on base change. The majority of base changes are T>C (A>G) and G>A (C>T). **d**, Mutational spectrum of all SNVs (top) and low-frequency SNVs (NHL < 1%, bottom) detected in mouse oocytes. The oocytes show a similar pattern of mutational spectrum within the strain, while different strains are significantly different in mutational spectrum. **e**, Heatmaps of clustering of individual mtDNA detected in B6 mouse oocytes based on SNVs (top panels: NHL < 1%, bottom panels: all SNVs).

**Supplementary Fig. 5.**
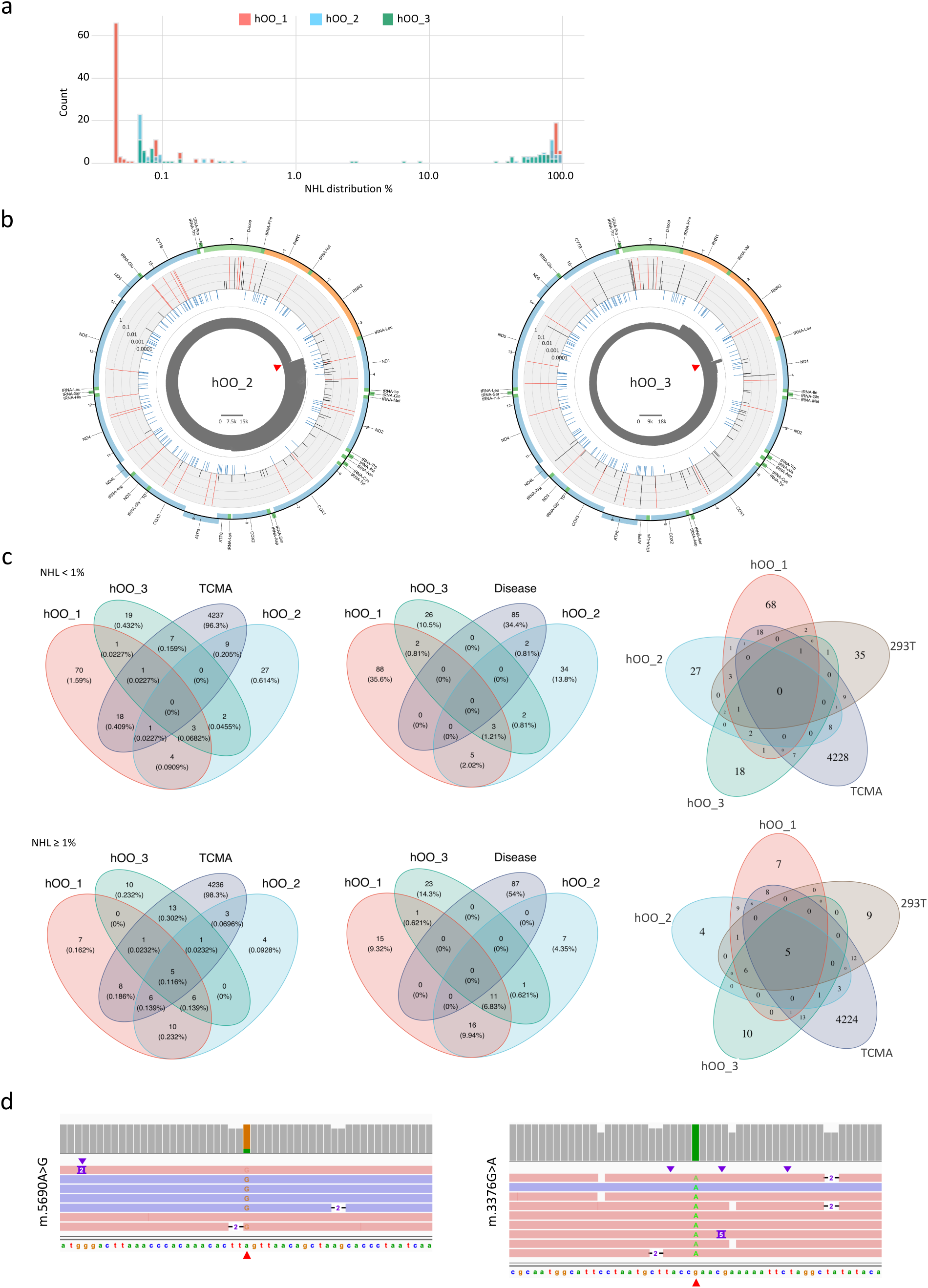
SNV analysis in the iMiGseq of human oocytes. **a**, Distribution of NHL of mtDNA SNVs detected in the three hOOs. The SNVs fall into two groups, the low-frequency SNVs have NHLs below 1%, while the high-frequency ones have NHLs above 30%. **b**, Circular plots showing the distribution of SNVs in the mitochondrial genome of hOO_2 and hOO_3. The arrangement of the circular plots is similar to Fig. 3a. **c**, Venn diagrams similar to those shown in Fig. 3f, except that the variants are divided into two categories. The upper panels show SNVs with NHL < 1%, and the lower panels are for NHL ≥ 1%. **d**, The two disease related SNVs detected in hOO_2 shown in IGV tracks. The mutant bases are indicated by red triangles under the plots.

**Supplementary Fig. 6.**
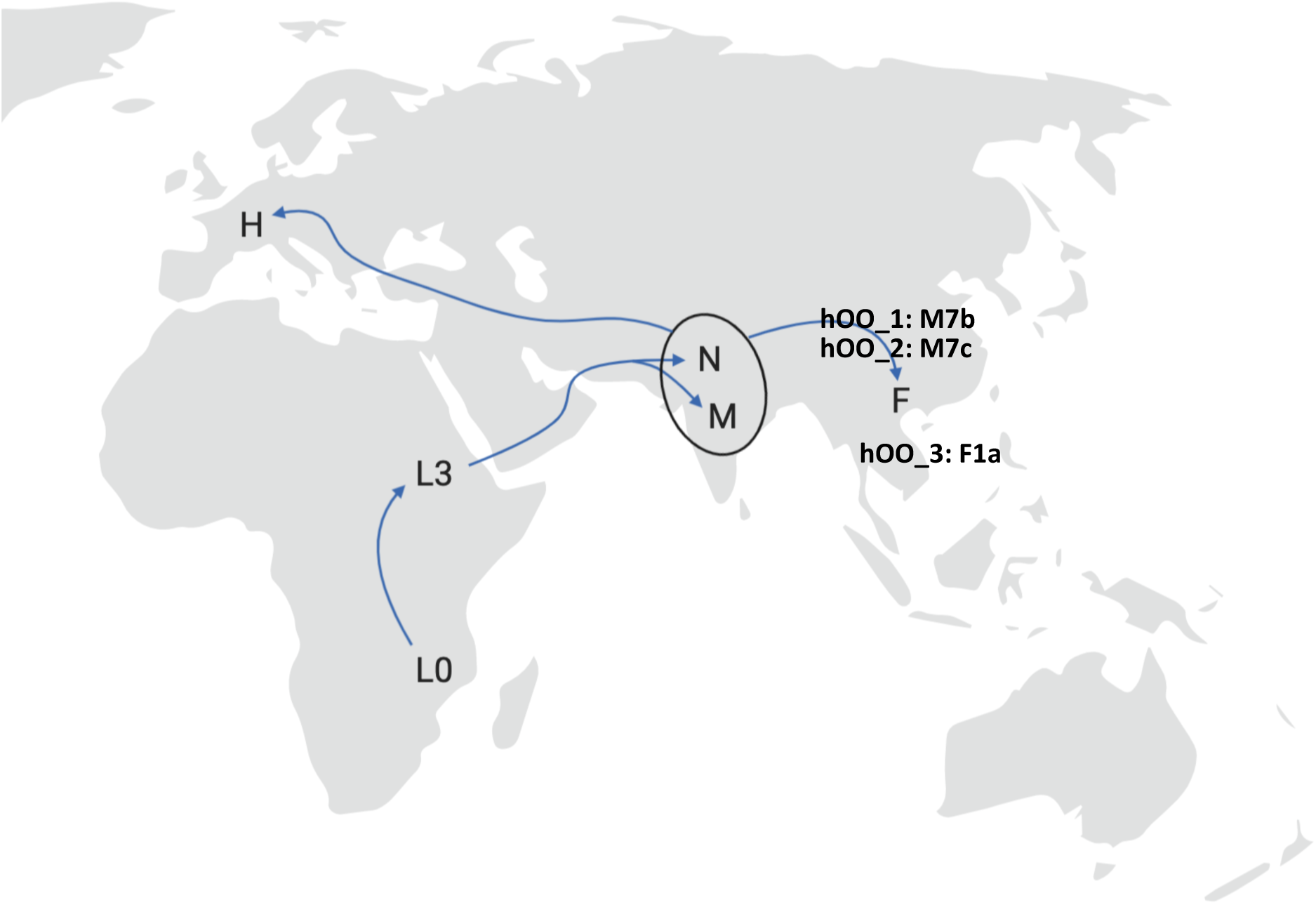
A map showing the mtDNA phylogeny and geographic distribution of the haplogroups of the three hOOs in this study.

**Supplementary Fig. 7.**
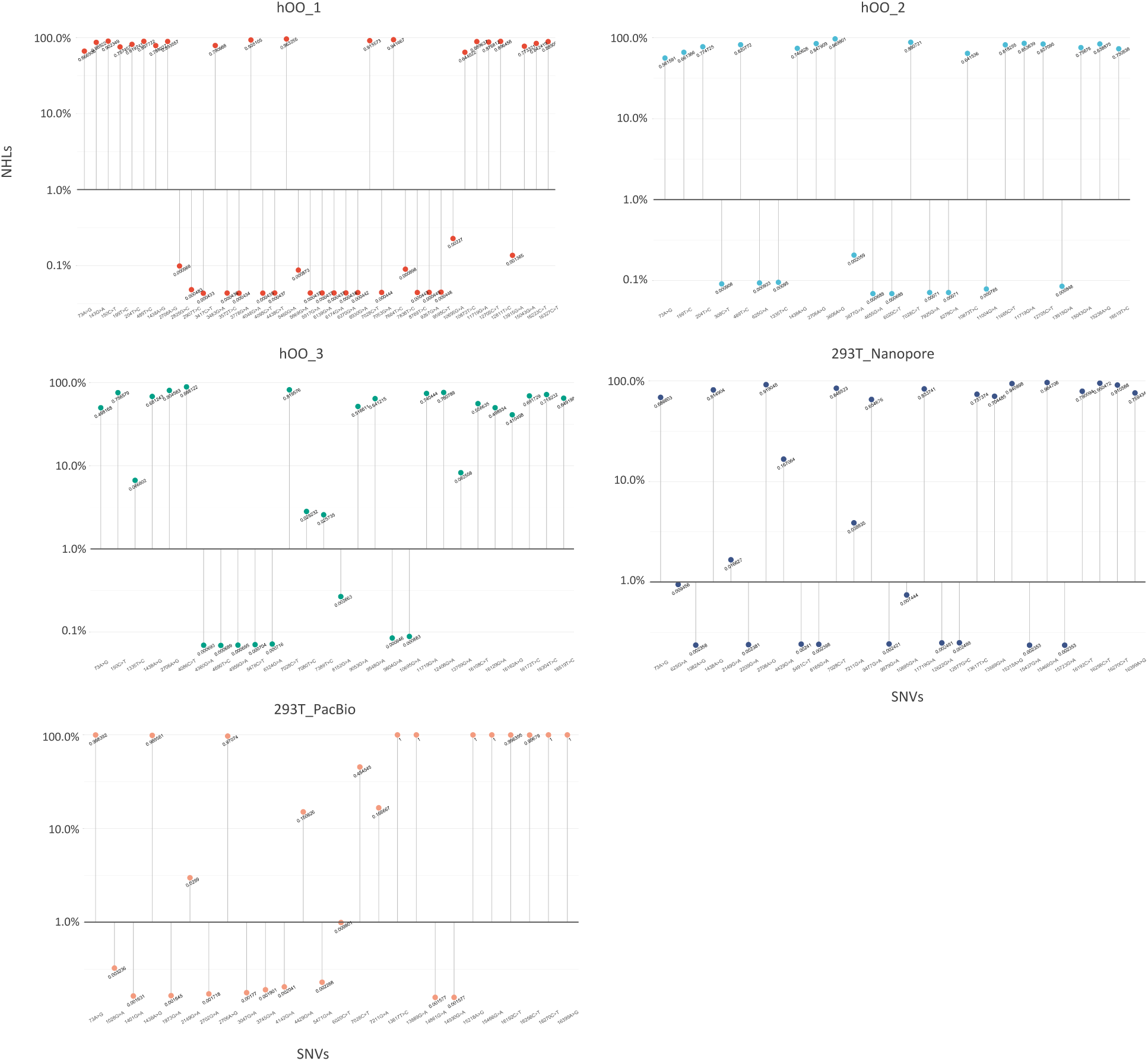
VAF of SNVs overlapping with TCMA SNVs in human oocytes and 293T cells. A significant number of cancer related SNVs exist below the 1% detection limit of current methods.

**Supplementary Fig. 8.**
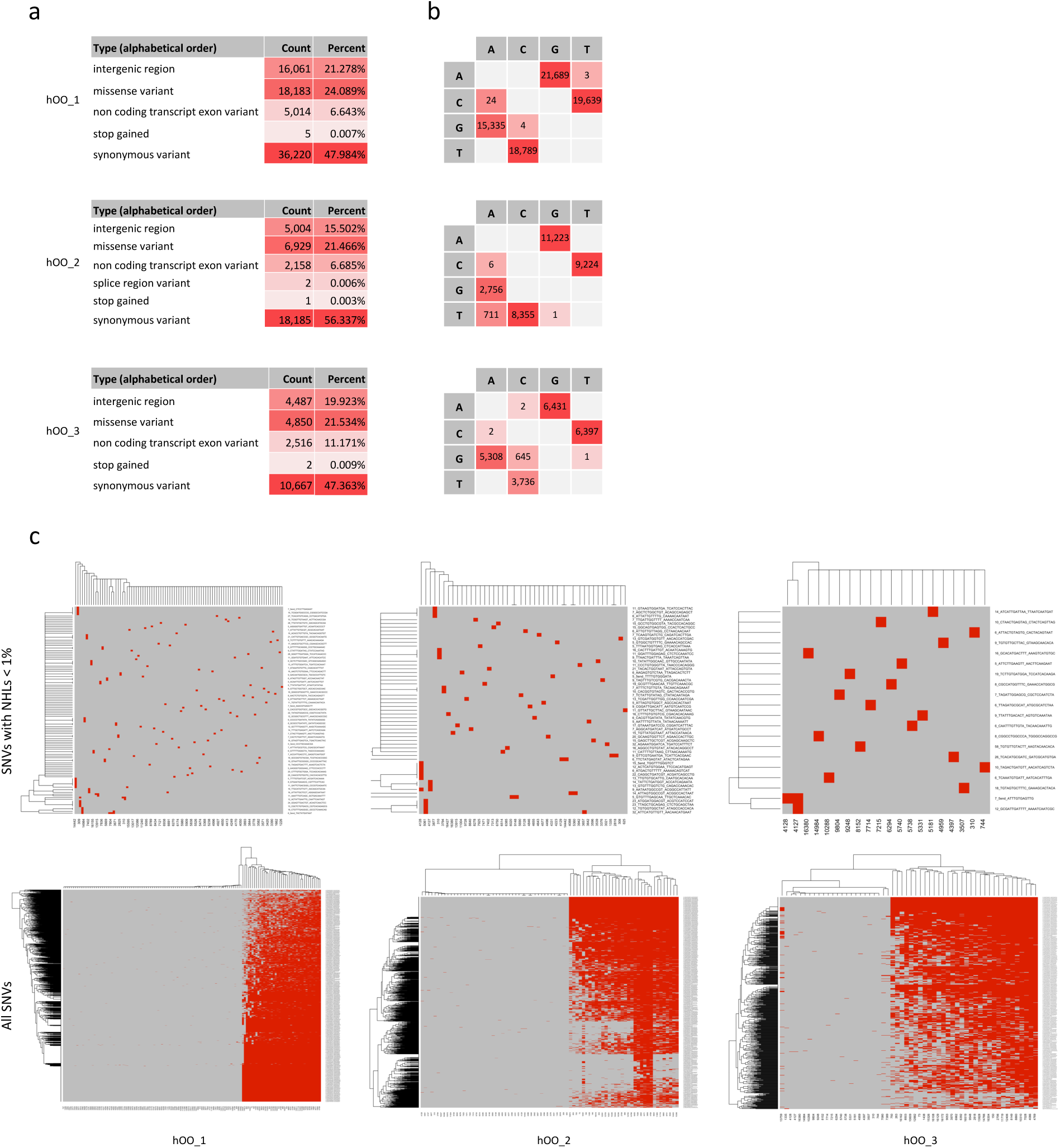
SNV annotation for human oocytes. **a**, Analysis of mtDNA SNVs of hOOs based on functional annotation. The majority of SNVs are synonymous variants in all oocytes. **b**, Analysis of mtDNA SNVs of hOOs based on base change. The majority of base changes are A>G (T>C) and C>T (G>A). **c**, Heatmaps of clustering of individual mtDNA detected in hOOs based on mtDNA SNVs (top panels: NHL < 1%, bottom panels: all SNVs).

**Supplementary Fig. 9.**
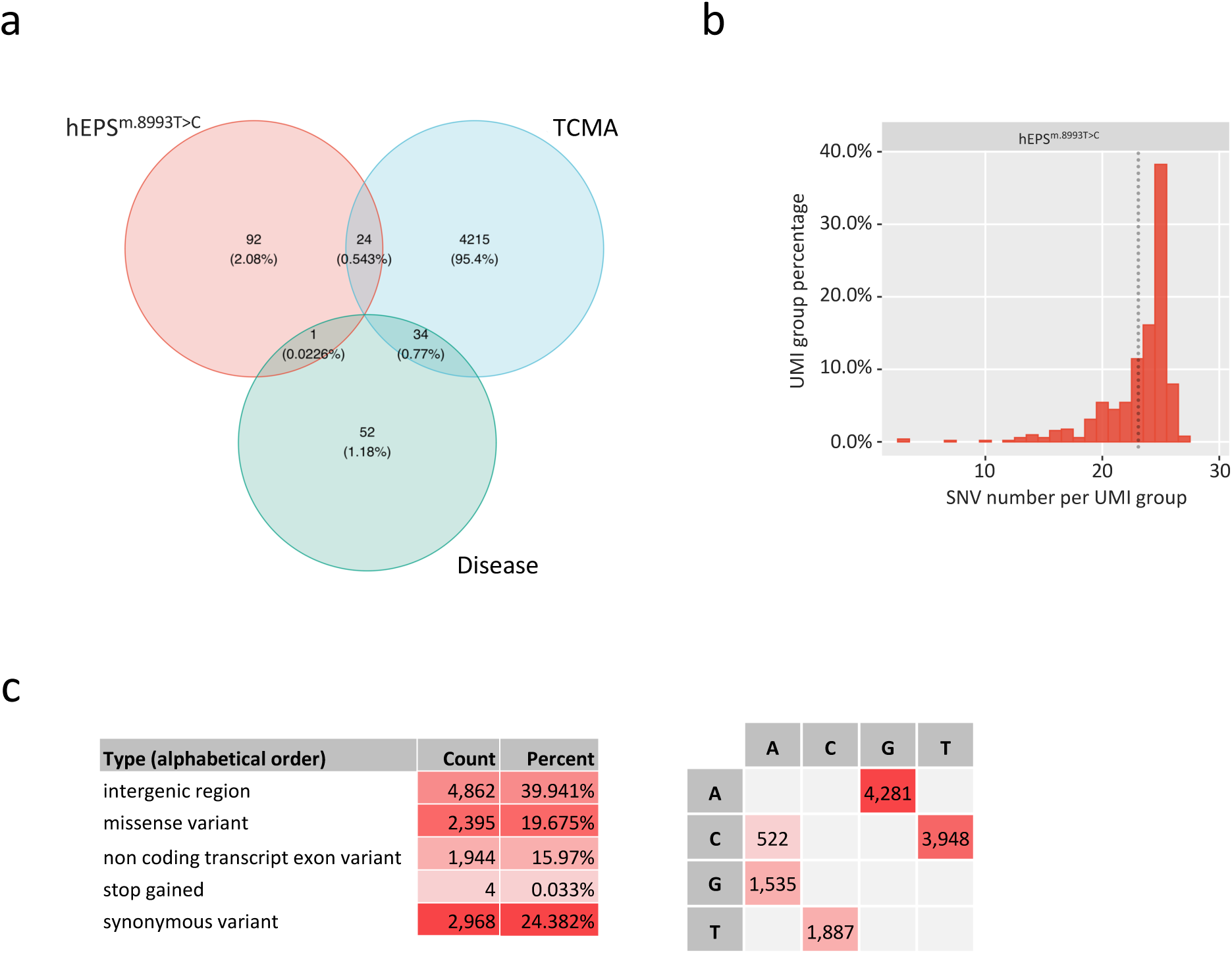
iMiGseq of patient derived hEPS^m.8993T>C^ cells. **a**, A Venn diagram showing the overlap of mtDNA SNVs in hEPS^m.8993T>C^, cancers (TCMA) and confirmed pathogenic SNVs (Disease). The m.8993T>C mutation exists in both hEPS cells and the Disease database. **b**, Histogram of SNV load per mtDNA in hEPS^m.8993T>C^ cells. The median SNV number is indicated by the vertical dotted line. **c**, Analysis of mtDNA SNVs of hEPS^m.8993T>C^ based on functional annotation and base change.

**Supplementary Table 1.** The distribution of SNVs (NHL < 1%) among mtDNA regions in hOOs.

| Region | bp | Expected % | hOO-1 |  |  | hOO-2 |  |  | hOO-3 |  |  |
| --- | --- | --- | --- | --- | --- | --- | --- | --- | --- | --- | --- |
|  |  |  | Expected | Observed | p-value<sup>‡</sup> | Expected | Observed | p-value<sup>‡</sup> | Expected | Observed | p-value<sup>‡</sup> |
| D-loop | 1122 | 6.8 % | 8.9 | 10 | 0.84* | 4.0 | 6 | 0.44* | 2.4 | 2 | 0.94* |
| tRNA | 1508 | 9.1 % | 11.9 | 10 | 0.67 | 5.4 | 3 | 0.40* | 3.2 | 2 | 0.69* |
| rRNA | 2513 | 15.2% | 19.9 | 5 | **0.00045** | 9.0 | 3 | **0.047\*** | 5.3 | 2 | 0.18* |
| Synonymous | 2834 | 17.1% | 22.4 | 35 | **0.005** | 10.1 | 19 | **0.0036** | 6.0 | 11 | **0.043\*** |
| Non-synonymous | 8533 | 51.5% | 67.5 | 71 | 0.60 | 30.4 | 28 | 0.62 | 18.0 | 18 | 0.87 |
| Total | 16569<sup>#</sup> |  |  | 131<sup>†</sup> |  |  | 59<sup>†</sup> |  |  | 35<sup>†</sup> |  |

